# PI3Kα inhibition blocks osteochondroprogenitor specification and the hyper-inflammatory response to prevent heterotopic ossification

**DOI:** 10.1101/2023.09.14.557704

**Authors:** José Antonio Valer, Alexandre Deber, Marius Wits, Carolina Pimenta-Lopes, Marie-José Goumans, José Luis Rosa, Gonzalo Sánchez-Duffhues, Francesc Ventura

**Affiliations:** Departament de Ciències Fisiològiques, Universitat de Barcelona, IDIBELL, C/ Feixa Llarga s/n 08907 Hospitalet de Llobregat, SPAIN; Department of Cell and Chemical Biology, Leiden University Medical Center, Einthovenweg 20, 2333 ZC, Leiden, the Netherlands; Nanomaterials and Nanotechnology Research Center (CINN-CSIC), Health Research Institute of Asturias (ISPA), 33011 Oviedo, Asturias, Spain

**Keywords:** bone/ heterotopic ossification/fibrodysplasia ossificans progressiva/ BMP/ activin A/ PI3K/ Alpelisib/ BYL719/ inflammation/ muscle regeneration

## Abstract

Heterotopic ossification (HO) occurs following mechanical trauma and burns, or congenitally in patients suffering from fibrodysplasia ossificans progressiva (FOP). Recently, we demonstrated that inhibitors of phosphatidyl-inositol 3-kinase alpha (PI3Kα) may be a useful therapy for patients undergoing HO. In this study, using the already marketed BYL719/Alpelisib/Piqray drug, we have further confirmed these results, detailed the underlying mechanisms of action, and optimized the timing of the administration of BYL719. We found that BYL719 effectively prevents HO even when administered up to three to seven days after injury. We demonstrate in cell cultures and in a mouse model of HO that the major actions of BYL719 are on-target effects through the inhibition of PI3Kα, without directly affecting ACVR1 or FOP-inducing ACVR1^R206H^ kinase activities. *In vivo*, we found that a lack of PI3Kα in progenitors at injury sites is sufficient to prevent HO. Moreover, time course assays in HO lesions demonstrate that BYL719 not only blocks osteochondroprogenitor specification, but also reduces the inflammatory response. BYL719 inhibits the migration, proliferation and expression of pro-inflammatory cytokines in monocytes and mast cells, suggesting that BYL719 hampers the hyper-inflammatory status of HO lesions. Altogether, these results highlight the potential of PI3Kα inhibition as a safe and effective therapeutic strategy for HO.

## Introduction

Heterotopic ossification (HO) is a disorder characterized by ectopic bone formation at extraskeletal sites, including skeletal muscle and connective tissues. Trauma-induced HO develops as a common post-operative complication after orthopedic surgeries (e.g., hip arthroplasty), blast injuries, deep burns and nervous system injuries (Hwang et al., 2022). In addition to trauma-induced HO, fibrodysplasia ossificans progressiva (FOP) is a rare congenital autosomal dominant disorder also involving HO (Bravenboer et al., 2015). Ectopic bones form progressively through endochondral ossification, mostly in episodic flare-ups associated with inflammation (Towler and Shore, 2022). These cumulative osteogenic events lead to reduced mobility resulting from ankylosing joints. Affected individuals have a shorter life-span, most commonly due to thoracic insufficiency syndrome (Kaplan et al., 2010; Pignolo et al., 2016).

HO reflects a shift from a normal tissue repair process to an aberrant reactivation of bone-forming developmental programs that require acute inflammation and the excessive expansion of local progenitors, followed by inappropriate differentiation of these progenitors into chondroblasts and osteoblasts which finally results in bone formation (Hwang et al., 2022). In both trauma-induced HO and FOP, pathology appears following the excessive activation of receptors sensitive to Transforming growth factor-β (TGFβ) superfamily members (Lees-Shepard et al., 2018; Wang et al., 2018). FOP arises from gain-of-function mutations in the bone morphogenetic protein (BMP) type I receptor *ACVR1*, with the most common mutation being c.617G>A, R206H (Shore et al., 2006). Higher SMAD1/5 mediated signaling of mutated ACVR1 has been partially attributed to a loss of auto-inhibition of the receptor and mild hypersensitivity to BMP ligands (Chaikuad et al., 2012; Groppe et al., 2011; Van Dinther et al., 2010). Signaling by mutated ACVR1 still relies on ligand-induced heterotetrameric clustering but, unlike wild type ACVR1, does not require ACVR2A/B kinase activity (Agnew et al., 2021; Ramachandran et al., 2021). More importantly, mutated ACVR1 receptors alter their signaling specificity, abnormally activating SMAD1/5 and altering the non-canonical BMP signals (e.g., p38 or PI3K) in response to activin A (Hatsell et al., 2015; Hino et al., 2015; Valer et al., 2019a). Accordingly, evidence supports that activin A is necessary and sufficient for HO in FOP (Hatsell et al., 2015; Lees-Shepard et al., 2018; Upadhyay et al., 2017). In contrast, activin A does not drive post-traumatic HO (Hwang et al., 2020), and non-genetically-driven HO mostly arises from excessive BMP and TGFβ signaling, with functional redundancy between different type I receptors of the TGFβ-superfamily (Agarwal et al., 2017; Patel et al., 2022; Sorkin et al., 2020; Wang et al., 2018).

Evidence points to mesenchymal fibroadipogenic precursors (FAPs), which are widely distributed in muscle and other connective tissues, as the key cell-of-origin that aberrantly undergo chondrogenesis and further ectopic bone formation (Dey et al., 2016; Eisner et al., 2020; Lees-Shepard et al., 2018). However, both FOP or non-genetic HO also require local tissue destruction and inflammation, which indicate that FAPs should be invariably primed for ectopic bone formation by this inflammatory microenvironment (Barruet et al., 2018; Convente et al., 2018; Hwang et al., 2022; Matsuo et al., 2019; Pignolo et al., 2013; Sorkin et al., 2020).

Inflammation in HO is characterized by an initial acute response following injury with the activation of innate immunity and the influx of neutrophils and monocytes (Convente et al., 2018; Hwang et al., 2022). Additionally, B and T cells of the adaptive system are also recruited (Chakkalakal et al., 2012; Convente et al., 2018). However, mice lacking B or T lymphocytes exhibited no delay in the development of heterotopic ossification (HO) after injury, indicating that these cells may play a subsequent role in the dissemination of bone lesions (Kan et al., 2009). During intermediate and late inflammatory stages, the recruitment of monocytes, macrophages and mast cells occurs, which in turn exerts autocrine and paracrine effects on nearby FAPs (Chakkalakal et al., 2012; Convente et al., 2018; Sorkin et al., 2020; Tu et al., 2022). Highlighting their relevance in HO, the depletion of mast cells and macrophages profoundly impairs genetic and non-genetic HO (Convente et al., 2018; Torossian et al., 2017; Tu et al., 2022). BMP receptors are robustly expressed in monocytes and macrophages and the expression of *ACVR1^R206H^* has been shown to extend inflammatory responses in patient-derived macrophages (Barruet et al., 2018; Matsuo et al., 2021, 2019). Among the plethora of cytokines secreted by monocytes, macrophages and mast cells, both activin A and TGF-β stand out as extremely relevant for HO (Alessi Wolken et al., 2017; Hatsell et al., 2015; Lees-Shepard et al., 2018; Patel et al., 2022; Sorkin et al., 2020; Upadhyay et al., 2017).

Genetic and pharmacological studies have indicated that osteochondroprogenitor specification and maturation depend on phosphatidylinositol 3-kinase-α (PI3Kα) (Ford-Hutchinson et al., 2007; Fujita et al., 2004; Gámez et al., 2016; Ikegami et al., 2011). We found that PI3K signaling was also linked to HO, since inhibitors of PI3Kα (BYL719/Alpelisib/Piqray) prevented HO in mouse models without major side-effects (Valer et al., 2019a). Mechanistically, PI3Kα inhibitors hamper canonical and non-canonical BMP signaling, decreasing total and phosphorylated SMAD1/5 levels and reducing transcriptional responsiveness to BMPs/Activin A in mesenchymal progenitors (Gámez et al., 2016; Valer et al., 2019b). In the current study, we demonstrate that the pharmacological and genetic inhibition of PI3Kα in HO progenitors at injury sites reduces HO *in vivo*. Moreover, envisioning a future translation into the clinic, we have optimized the administration of BYL719 and found that the delayed administration of BYL719, up to seven days after inflammatory injury, still prevents HO in mice. In addition, we found that BYL719 blocks osteochondroprogenitor specification and effectively reduces the essential inflammatory response. Altogether, the data presented here show the potent therapeutic effect of PI3Kα inhibition on HO in mouse models.

## Results

### Delayed initiation of treatment with PI3K***α*** inhibitor effectively prevents heterotopic ossification

Heterotopic ossification takes place following a precise temporal pattern of progenitor activation and the recruitment of distinct cell types at the ossification centers which causes early divergence from the normal skeletal muscle repair program. We previously found that the pharmacological administration of BYL719 prevents HO in a mouse model of HO (Valer et al., 2019a). Given that BYL719 is marketed for the treatment of cancer and overgrowth syndrome, it is a promising therapeutic molecule in pathological HO. To further determine a potential therapeutic window for BYL719 and better understand the cellular targets of BYL719, we evaluated the effect of BYL719 when administered intermittently and several days after HO induction. For this, we used a conditional mouse model (*ACVR1^Q207Dfl/fl^)* in which we combine the overexpression of Cre recombinase through adenoviral particles with the injection of cardiotoxin intramuscularly in the hindlimb (Fukuda et al., 2006). In this model, we administered intermittently BYL719 (i.p. 25 mg/kg) or vehicle control starting one, three or seven days after HO induction. We also implemented a different administration regimen of BYL719 only for the initial three days following the injury (Figure 1A). None of the treatments led to significant changes in mouse body weight (Figure 1-Figure Supplement 1). HO volume was analyzed by micro computed tomography (µCT) twenty-three days post-injury. The early administration of the treatment with BYL719 at day one or three after the onset of HO resulted in significantly lower HO; late administration seven days after the induction of HO was also partially effective (Figure 1B and C). Discontinuation of the treatment after three days did not prevent HO at day 23. These results indicate that the inhibition of PI3Kα by BYL719 after injury prevents HO and this protection is still effective if the treatment is started several days after injury, expanding its therapeutic window.

**Figure 1.**
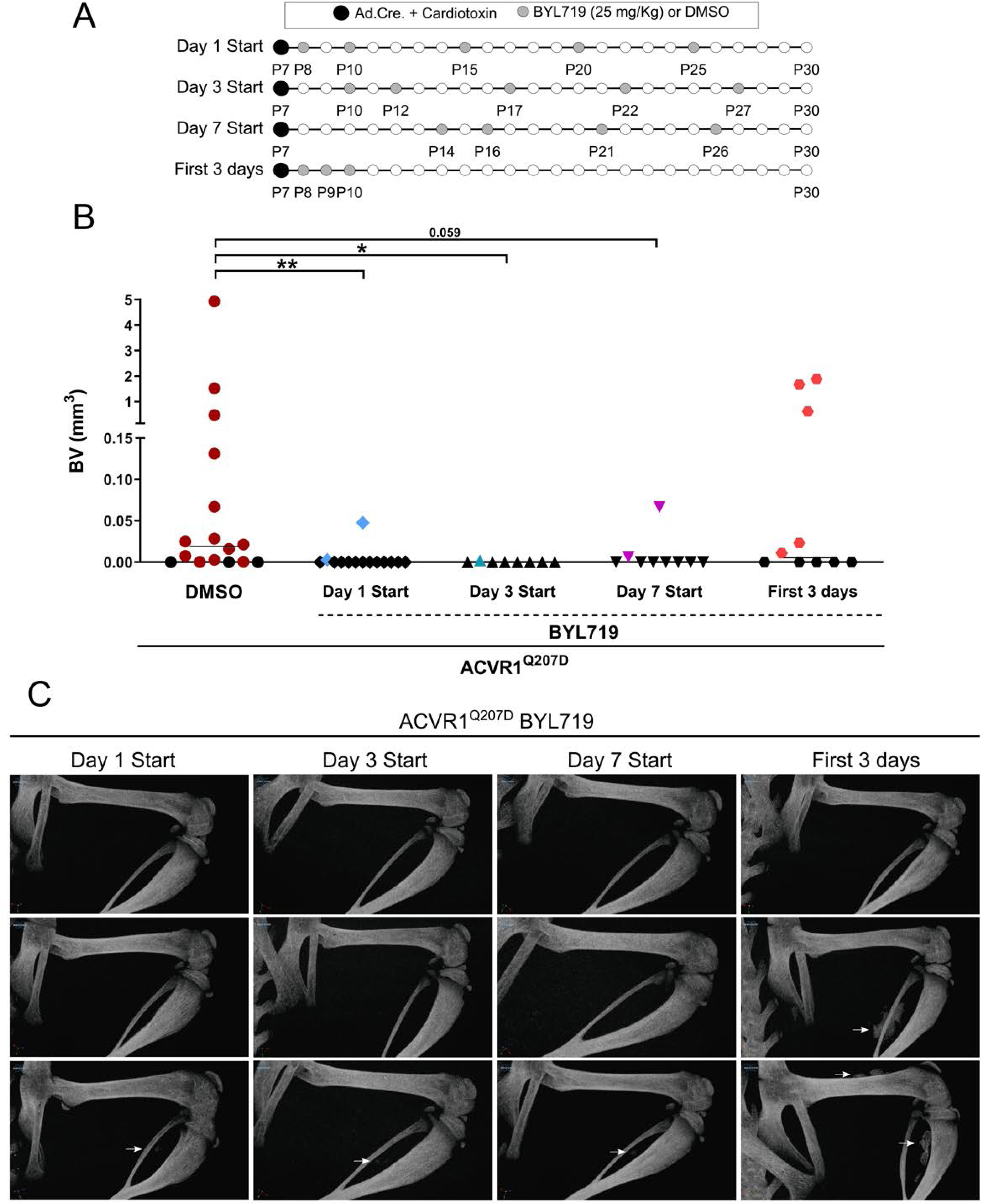
(A) Heterotopic ossification was induced at P7 through the injection of adenovirus-Cre (Ad.Cre) and cardiotoxin in the mice hindlimb. The pattern of 5 administrations of DMSO or BYL719 (25 mg/kg) is indicated by grey dots started at P8 (start day 1), P10 (delayed start day 3), and P14 (delayed start day 7). One group was administered DMSO or BYL719 (25 mg/kg) only at P8, P9, and P10 (first 3 days post-HO-induction), as indicated by grey dots. The final time-point, P30, was conserved between experimental groups. (B) Quantification of heterotopic ossifications bone volume (BV) (mm^3^) of each experimental group. Colored symbols indicate the presence of heterotopic ossifications. Black symbols indicate the absence of heterotopic ossifications. Individual mouse values with group median are shown. *P<0.05, **P<0.01, Kruskal–Wallis test with Dunn’s multiple comparison test. (C) Representative 3D microtomography images of the injected hindlimbs of 3 different mice for each experimental group. White arrows point to heterotopic ossification.

### Deficiency of PI3K***α*** at injury sites is sufficient to partially prevent heterotopic ossification

Since other small molecules designed to target PI3Kα have reported partial off-target effects on kinases other than PI3Kα (Furet et al., 2013; Jamieson et al., 2011), we aimed to confirm that the results obtained with BYL719 were due to the specific inhibition of PI3Kα. Therefore, we developed a conditional mouse model (*ACVR1^Q207Dfl/fl^:p110*α*^fl/fl^*) in which the Cre recombinase simultaneously drives the expression of *ACVR1^Q207D^*and the deletion of the catalytic subunit of PI3Kα (*p110*α) upon intramuscular hindlimb combined injection of adenovirus-Cre and cardiotoxin to trigger HO. With this approach, Cre mediates the expression of ACVR1 *^Q207D^* and the deletion of PI3Kα in the same cells, whereas cells recruited afterwards will remain wild type for both genes. As expected, µCT analysis performed 23 days after injury showed extensive HO in mice expressing ACVR1^Q207D^, which was prevented by intermittent treatment with BYL719 initiated one day after injury (Figure 2A, 2B and 2D). Genetic deletion of *p110*α in mice expressing ACVR1^Q207D^ led to a reduction in HO, which was further enhanced by BYL719 (Figure 2B). Importantly, no significant changes in the weight of the mice were observed, confirming the absence of severe toxicity (Figure 2-Figure Supplement 1A). Histomorphometrically, ACVR1^Q207D^ mice wild type for PI3Kα developed islands and/or bone spurs with a well-organized bone structure. However, in PI3Kα deficient mice ACVR1^Q207D^ expression only led to minor ectopic calcifications that were already surrounded by fully regenerated muscle tissue on the 23^rd^ day after injury (Figure 2D, Figure 2-Figure Supplement 1B, Figure 2-Figure Supplement 2). Accordingly, analysis of the ratio bone volume/tissue volume (BV/TV) and the correlation between bone volume and BV/TV of individual mice was clearly different between *p110*α genotypes, irrespective of BYL719 treatment (Figure 2C, Figure 2-Figure Supplement 1C). These results demonstrate that the abrogation of PI3Kα activity (either genetic of pharmacologically) in cells at injury sites is relevant for their inhibitory effects in HO. This might be attributed to the direct effects of BYL719 on progenitor cell expansion and chondroblast specification, as well as its possible ability to reduce the inflammatory response required for HO.

**Figure 2.**
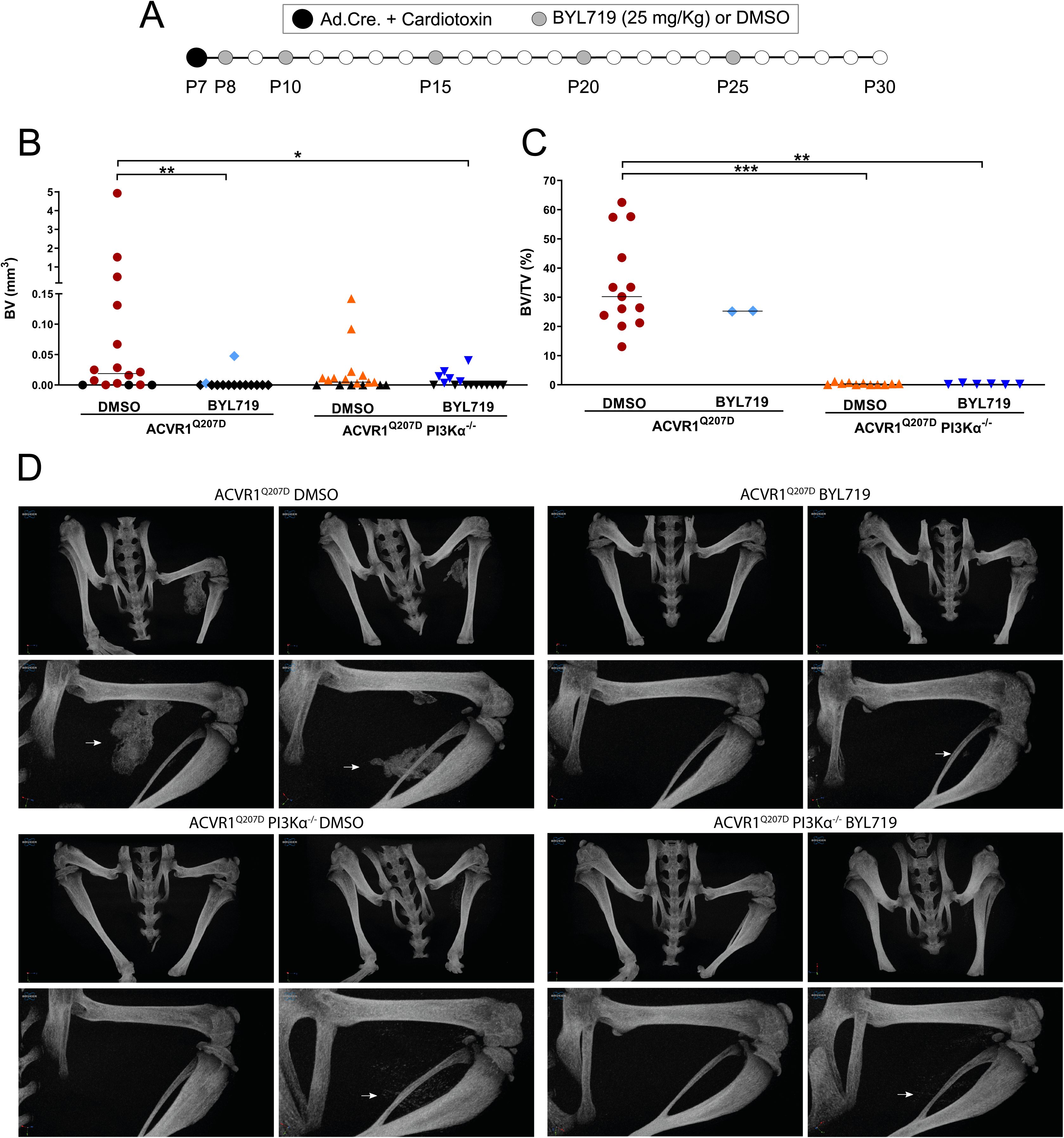
(A) Heterotopic ossification was induced at P7 through the injection of Adenovirus-Cre (Ad.Cre) and cardiotoxin in the mice hindlimb. Either DMSO (vehicle) or BYL719 (25 mg/kg) were injected following the scheme indicated with grey dots, starting at P8 (B) Quantification of heterotopic ossifications bone volume (BV)(mm^3^) of each experimental group. Colored symbols indicate the presence of heterotopic ossifications. Black symbols indicate the absence of heterotopic ossifications. Individual mouse values with group median are shown. *P<0.05, **P<0.01, Kruskal–Wallis test with Dunn’s multiple comparison test. (C) Quantification of the ratio of bone volume per tissue volume (bone volume/tissue volume, BV/TV) within heterotopic ossifications of each experimental group, including only mice with detected HO. Data shown are of each individual mouse with the group median. **P<0.01, ***P<0.001, Kruskal-Wallis test with Dunn’s multiple comparison test. (D) Representative 3D frontal microtomography images of the injected hindlimbs of mice for each experimental group and a detailed close-up image for each selected mouse. White arrows show heterotopic ossification in the close-up images.

### Genetic or pharmacological inhibition of PI3K***α*** prevents osteochondroprogenitor specification and reduces the number of FAPs at injury sites

Next, we aimed to assess the effects of PI3Kα inhibition in osteochondroprogenitor cells. For this, we isolated bone marrow-derived mesenchymal stem cells (BM-MSCs) from *p110*α*^fl/fl^* mice and transduced them *ex-vivo* with retrovirus expressing *Acvr1* (*wild type* and *R206H*) with or without Cre recombinase. The gene expression levels of exogenous mRNA of *Acvr1* and *Acvr1^R206H^* was similar (Figure 3A). As anticipated, transduction with Cre viruses significantly reduced the expression levels of p110α to approximately 40%. This is consistent with an estimated transduction efficiency of nearly 60% (Figure 3B). The overexpression of *Acvr1^R206H^* increased basal and activin-dependent expression of canonical (*Id1* and *Sp7*) and activin-dependent expression of non-canonical (*Ptgs2*) BMP target genes (Figure 3C), known markers and drivers of osteochondrogenic differentiation (Nakashima et al., 2002; Wang et al., 2013). Both pharmacological and genetic inhibition of PI3Kα reduced canonical and non-canonical ACVR1^R206H^ transcriptional responses induced by activin A. Of note, stimulation with activin A increased its own transcription (*Inhba*) in *Acvr1^R206H^*-transduced BM-MSCs, which was also inhibited by BYL719 (Figure 3C). Therefore, BYL719 may also prevent osteochondroprogenitor differentiation indirectly, via transcriptional inhibition of *INHBA*/Activin A expression. These results demonstrate a direct effect of BYL719 on osteochondroprogenitor cells and suggest that the effects of BYL719 on ACVR1-downstream signaling are mainly on target, due to PI3Kα kinase inhibition.

**Figure 3.**
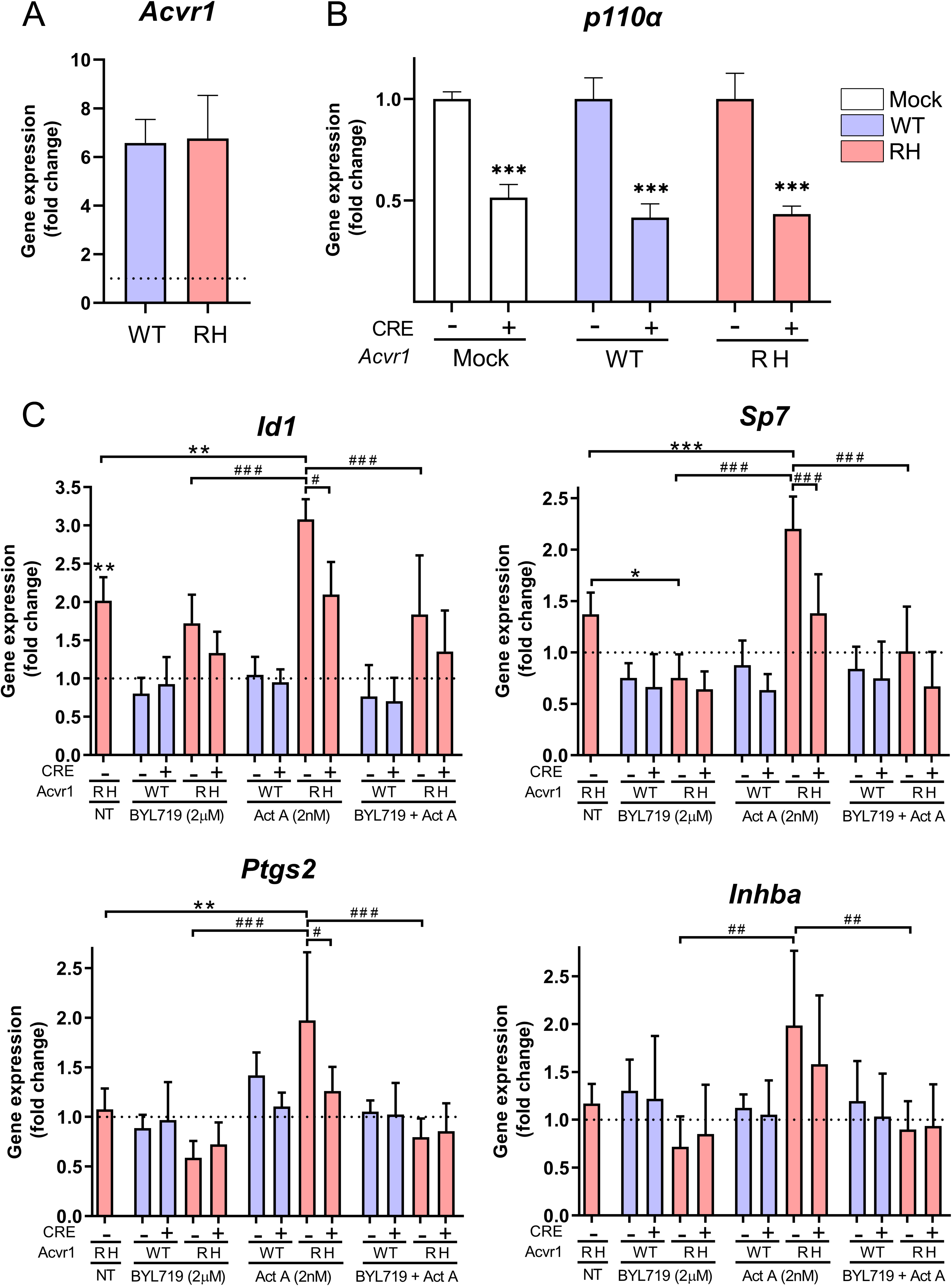
Analysis of *Acvr1* gene expression by qPCR in BM-MSCs from *p110* ^fl/fl^ mice infected with virus expressing wild-type *Acvr1* (WT) or *Acvr1^R206H^*(RH). (A) The endogenous expression level of *Acvr1* gene in mock-transfected BM-MSCs is shown as a dotted horizontal line. Data are shown as mean ± SD (n = 12 per group). Unpaired t-test between transfected groups. (B) Gene expression analysis of *p110*α in BM-MSCs from *p110α^fl/fl^* mice, infected with virus expressing wild-type *Acvr1* (WT) or *Acvr1*^R206H^ (RH) and/or Cre co-infection. Data are shown as mean ± SD (n = 3 per group). ***P<0.001, two-way ANOVA with Tukey’s multiple comparisons test. (C) mRNA expression of canonical (*Id1* and *Sp7*), non-canonical (*Ptgs2*) target genes, and activin A (*Inhba*) in BM-MSCs *p110* ^fl/fl^ transfected with *Acvr1* (wild type or R206H) with or without Cre recombinase. Cells were NT (not treated), or treated with BYL719 (2µM) and/or activin A (2nM). Expression data was normalized to those of control cells which were transfected only with *Acvr1* WT without any treatment, shown as a dotted horizontal line. Asterisks (*) refer to the differences between different conditions of *Acvr1* RH cells compared to control cells. Hash signs (#) refer to the differences between different conditions of *Acvr1* RH cells compared to *Acvr1* RH cells without Cre recombinase and treated with activin A. Data are shown as mean ± SD (n = 6 per group). * or # P<0.05, ** or ## P<0.01, *** or ### P<0.001, two-way ANOVA with Tukey’s multiple comparisons test.

In addition, we analyzed the number of fibroadipogenic precursors (FAPs) during the progression of ectopic bone formation in our *in vivo* mouse model of HO. Samples of injured muscles of the conditional mouse model (*ACVR1^Q207Dfl/fl^)* were subjected to immunofluorescence to identify the number of PDGFRA+ cells (FAPs) at 4, 9,16 and 23 days after induction of HO. The number of FAPs remained sustained until day 16^th^ after injury, decreasing afterwards (Figure 3-Figure Supplement 1A, B). More importantly, treatment with BYL719 reduced the number of PDGFRA+ cells throughout the ossification process. We also observed an increase in the diameter of myofibers in animals treated with BYL719 (Figure 3-Figure Supplement 1A, C). These results suggest an improved muscle regeneration in animals treated with BYL719 compared to those undergoing HO.

It is known that 2-aminothiazole-derived PI3Kα inhibitors could also inhibit ACVR1 kinase activity (Jamieson et al., 2011). Therefore, we investigated whether BYL719 effects could be explained by the reduction in ACVR1 kinase activity. For this, we made use of a kinase target engagement approach. Upon transient transfection of Nanoluciferase-tagged TGF-β receptors, we incubated the cells with BYL719 and/or an ATP-like tracer analogue. In the absence of an ATP competitor molecule, the tracer analogue and Nanoluciferase enzyme stay in close proximity, thereby allowing for bioluminescence resonance energy transfer (BRET) between the Nanoluciferase (energy donor, emitting at 460 nm) and the tracer analogue (energy acceptor) Once excited, the acceptor molecule emits fluorescence (610 nm) (Figure 4A and B). BYL719 incubated at 1 and 10 µM failed to significantly inhibit the acceptor emission of the BMP type I receptors ACVRL11, ACVR1, BMPR1A, ACVR1B, TGFBR1 and the mutant receptor ACVR1^R206H^ (Figure 4C), without significant changes in the protein levels of each receptor (Figure 4-Figure Supplement 1). In addition, we demonstrated that BYL719 does not target the activity of the type II kinase receptors ACVR2A, ACVR2B, BMPR2 and TGFBR2. We confirmed these results using *in vitro* kinase activity assays with recombinant ACVR1^R206H^. BYL719 exhibited no major effects on ACVR1^R206H^ activity up to 10µM (Figure 4D).

**Figure 4.**
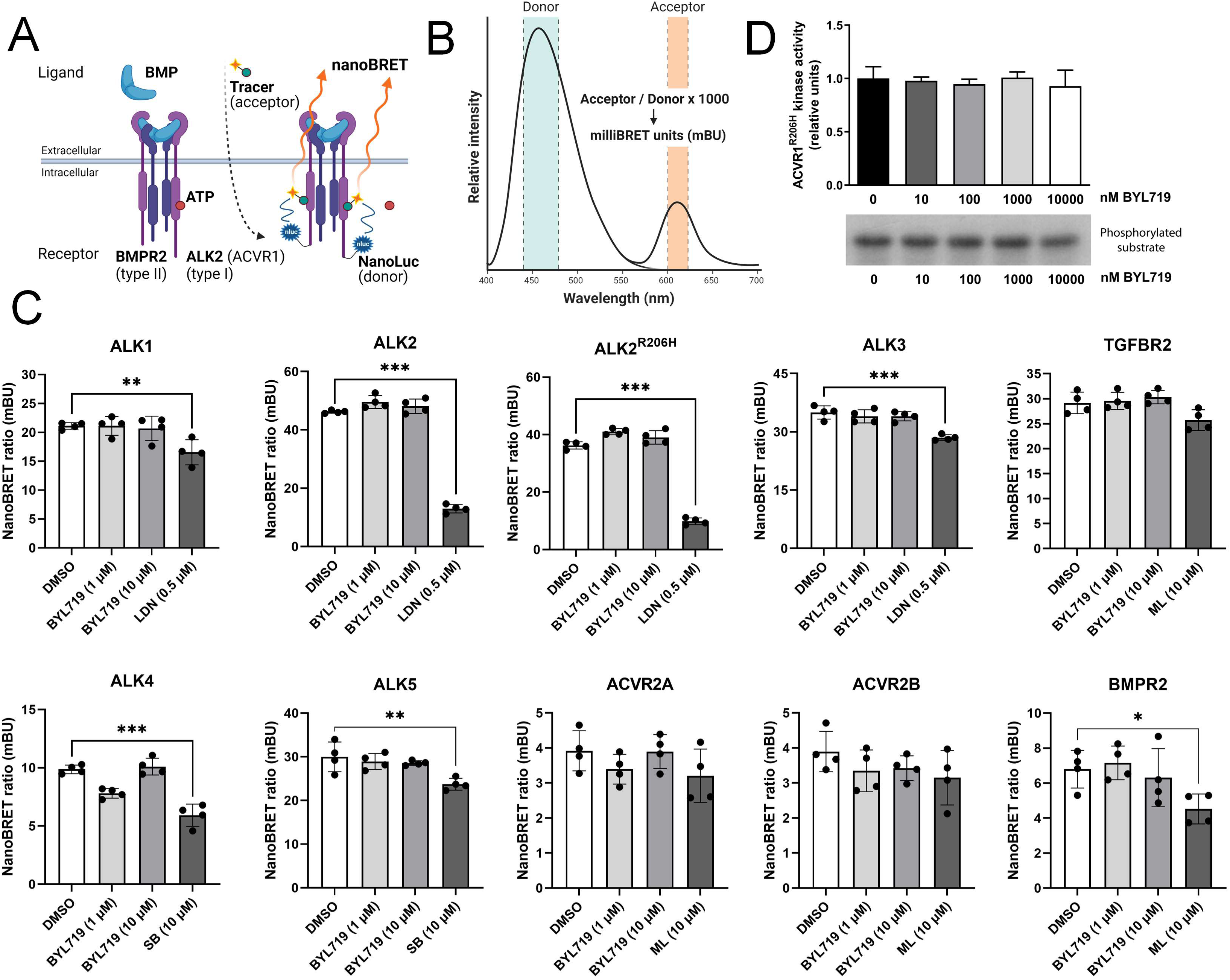
(A) A schematic depiction of nano-bioluminescence resonance energy transfer (NanoBRET) target engagement assays. TGF-β receptors-Nanoluciferase fusion proteins are expressed in COS-1 cells and an ATP-like tracer analogue in close proximity with the Nanoluciferase donor allows energy transfer from the Nanoluciferase donor to the fluorescent tracer acceptor. An ATP analogue molecule (type I inhibitor) will compete with the fluorescent tracer, impairing close proximity donor-acceptor and reducing the nanoBRET ratio. (B) The nanoBRET emission spectra consist of the Nanoluciferase donor (460nm) and the fluorescent acceptor (610nm). The nanoBRET ratio is shown as milliBRET units (mBU) by dividing the acceptor emission by the donor emission times 1000. (C) NanoBRET target engagement analyses of ACVRL1, ACVR1, ACVR1^R206H^, BMPR1A, ACVR1B, TGFBR1, TGFBR2, ACVR2A, ACVR2B and BMPR2 testing 1 or 10 µM BYL719 with n=4. As controls, LDN193189 (0,5 µM), SB431542 (10 µM), and ML347 (10 µM) were used. Data are shown as mean ± SD. One-way ANOVA with Dunnett’s multiple comparisons test. (D) Casein phosphorylation by ACVR1^R206H^ kinase. Phosphorylation was performed in the presence of ACVR1^R206H^ kinase and increasing concentrations of the PI3Kα inhibitor BYL719. Quantification of kinase activity. Data shown as mean ± SD (n=4 independent experiments). One-way ANOVA with Tukey’s multiple comparisons test. * P<0.05, ** P<0.01, *** P<0.001.

Given that BYL719 does not directly target the activity of ACVR1 or related receptors, we established two *in vitro* models to further profile the main targeted pathways. In human MSCs (hMSCs) stably overexpressing ACVR1 or ACVR1^R206H^, recombinant Activin A was able to induce SMAD1/5 activation and the expression of ACVR1 downstream targets *ID1* and *ID3* in cells over-expressing ACVR1^R206H^ (Figure 5-Figure Supplement 1A and B). We performed micromass chondrogenic differentiation assays in differentiation media supplemented with TGFβ1, with or without activin A and BYL719 in parental cells and cells transduced with ACVR1^WT^ or ACVR1^R206H^. Coincubation with BYL719 (10 μM) completely inhibited the formation of a chondrogenic glycosaminoglycans-rich matrix in response to TGFβ1 plus activin A in the different cell types (Figure 5A and B) and partially abrogated the expression of the chondrogenic genes *ACAN* and *MMP13* (Figure 5C). Furthermore, in these cells, coincubation with BYL719 (10 μM) was able to repress Activin A-induced alkaline phosphatase activity upon 7 and 11 days of culture, respectively (Figure 5D and E). In addition, to confirm these results, we isolated MSCs from UBC-CRE-ERT2/ACVR1^R206H^ ^fl/wt^ knock-in mice. These cells, when treated with 4OH-tamoxifen, express the intracellular exons of human *ACVR1^R206H^*in the murine *Acvr1* locus. Therefore, we can compare wild type and R206H MSCs isolated from the same mice. ACVR1^R206H^ mutant cells display an enhanced chondrogenic response to activin A compared to wild type cells (Figure 5F). In both wild type and mutant MSCs, treatment with BYL719 decreased the expression of chondrogenic. These results confirmed that BYL719 could inhibit chondrogenic differentiation of human and murine MSCs irrespective of the expression of either wild type or mutant ACVR1.

**Figure 5.**
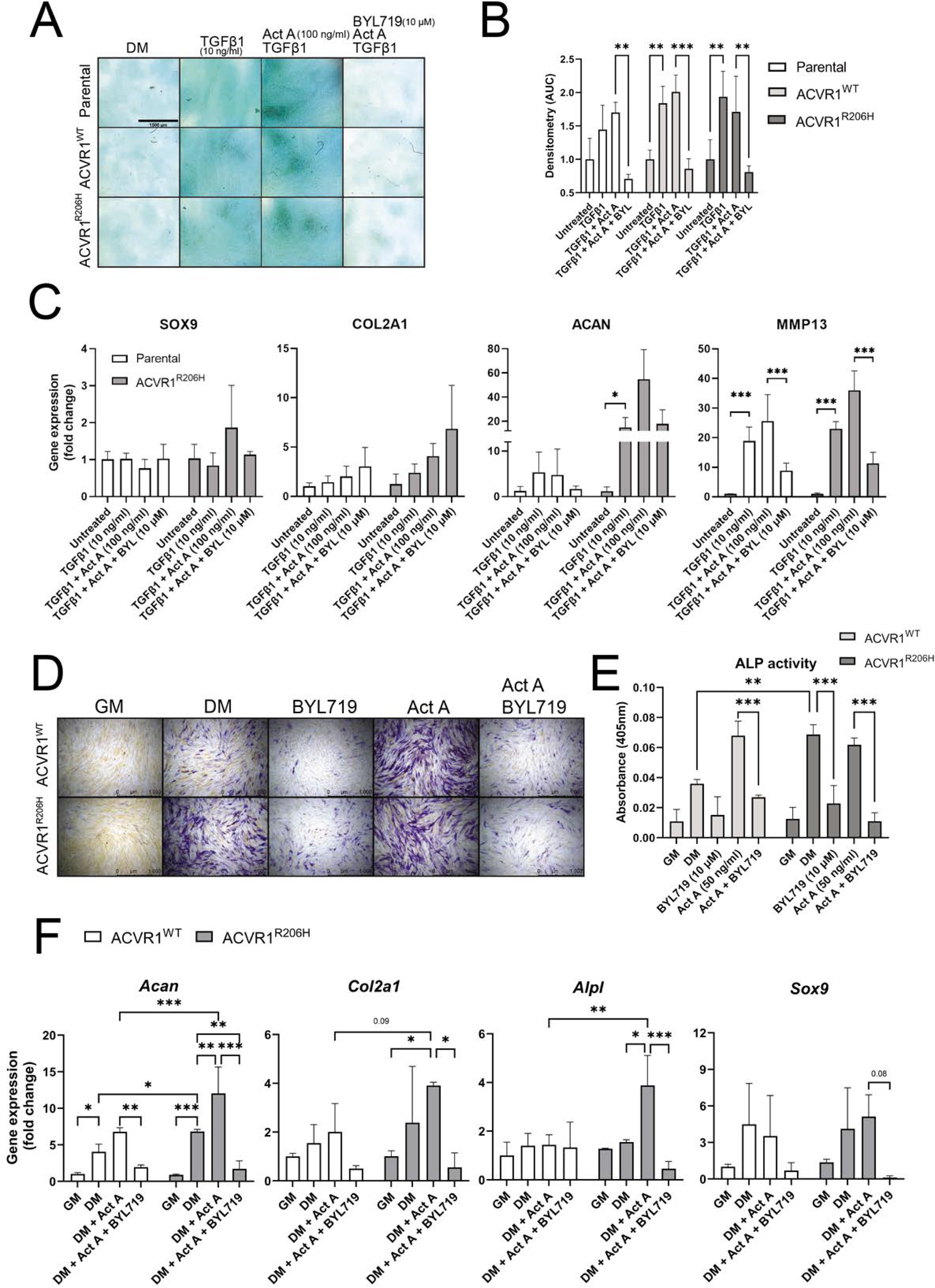
Analyses of chondrogenic progenitor specification of parental human MSCs (hMSCs), hMSC-ACVR1^WT^ or hMSC-ACVR1^R206H^ treated with TGF-β1 (10 ng/ml) and Activin A (100 ng/ml), with and without BYL719 (10 µM) as indicated (A) Representative images of Alcian Blue staining of micromass cultures after 3 weeks of chondrogenic differentiation. Scale bar represents 1500 µm. (B) Densitometry analysis of the Alcian Blue staining in (A) (n=3 per group). (C) RT-qPCR results of *SOX9, COL2A1, ACAN* and *MMP13* after chondrogenic progenitor specification (n=4 per condition). (D) Representative images of the ALP staining after one week of chondrogenic progenitor specification. Scale bar represents 1000 µm. (E) Quantification of the ALP activity (n=3 per group). (F) RT-qPCR results of *Acan, Col2a1, Alpl* and *Sox9* after chondrogenic progenitor specification from *Acvr1*^WT^ or ACVR1^R206H^ MSCs treated with Activin A (100 ng/ml), with or without BYL719 (10 µM) in TGFβ1-containing differentiation media (DM) as indicated (n=3 per condition). Data are shown as mean ± SD. In all figure panels, significance is detailed between the indicated conditions. * P<0.05, ** P<0.01, *** P<0.001, two-way ANOVA with Tukeýs multiple comparisons test.

To further identify the effects of BYL719, we performed bulk RNA sequencing (GSE237512) where we preincubated hMSCs-ACVR1^WT^ or hMSCs-ACVR1^R206H^ with or without BYL719 for 30 minutes, which was followed by stimulation with two high affinity ligands for ACVR1, Activin A (50ng/ml) or BMP6 (50ng/ml) for 1 hour (Figure 6A). Consistent with our results on osteogenic progenitor cells, gene ontology analysis of differentially expressed genes between cells expressing ACVR1^WT^ and ACVR1^R206H^ resulted in the robust regulation of relevant biological processes. Ossification (GO:0001503) and osteoblast differentiation (GO:0001649) were detected as two of the ten most significantly differentially regulated biological processes between these conditions (Figure 6B). Moreover, using gene ontology, we analyzed ossification and osteoblast differentiation biological processes in presence of ACVR1^WT^ or ACVR1^R206H^ receptor, with different ligands (BMP6 or Activin A), and with or without BYL719 inhibitor. The addition of BYL719 (1 µM) resulted in a downregulation of these GO terms for all conditions except activin A stimulated ACVR1^WT^ cells (Figure 6C).

**Figure 6.**
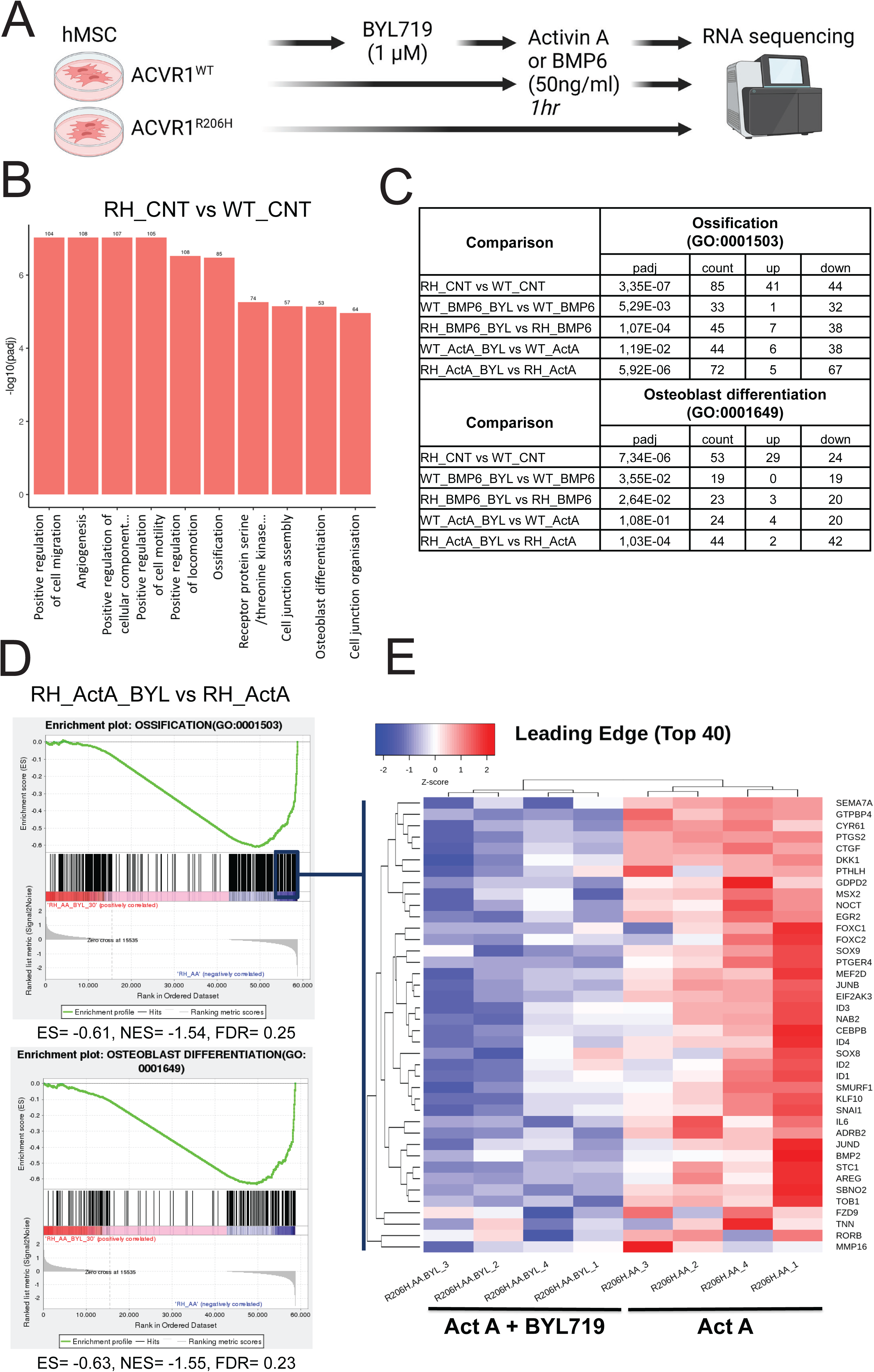
(A) Schematic depiction of the experimental setup, involving bulk RNA sequencing of human MSC-ACVR1^WT^ and ACVR1^R206H^ upon overnight starvation, 30 minutes pre-treatment with or without 1 µM BYL719 and with or without 1 hour Activin A (50 ng/ml) or BMP6 (50ng/ml) stimulation. (B) The top 10 most significant gene ontology (GO) terms using all (up- and down-regulated) differentially expressed genes between cells expressing ACVR1-^WT^ and ACVR1-^R206H^ under control conditions. Ossification (GO:0001503) and osteoblast differentiation (GO:0001649) were detected within the top 10 of the differentially regulated biological processes. (C) Table of GO terms ossification and osteoblast differentiation upon GO enrichment analysis in all tested conditions ACVR1^WT^ or ACVR1^R206H^ cells, stimulated with BMP6 or Activin A and treated with BYL719. Statistical significance upon GO enrichment analysis, count of total differentially expressed genes (DEGs), and their classification as up- or down-regulated DEGs are detailed for each comparison. D) Gene Set Enrichment Analysis (GSEA) with the groups RH_ActA_BYL vs RH_ActA, showing the enrichment plots of the gene ontology sets ossification (GO: 0001503) and osteoblast differentiation (GO:0001649). (E) A heatmap of the top 40 most relevant genes within the ossification GO geneset derived from the leading-edge subset of the enrichment plot (n=4 per group). Sample names are detailed as receptor (R206H) ligand (AA, Act A, Activin A)_inhibitor (BYL719, if present)_replicate#.

When comparing gene expressions profiles between ACVR1^R206H^ treated with Activin A, with or without BYL719, Gene Set Enrichment Analysis (GSEA) of our GO terms of interest showed negative normalized enrichment scores, consistent with our gene ontology analysis (Figure 6D). By looking at the leading edge we could investigate the most relevant genes within these GO terms, which included relevant genes of these pathways such as *PTGS2*, *MSX2*, *SOX9*, and *BMP2* (Figure 6E). Moreover, several inflammatory cytokine genes (e.g. *IL6* and *IL15*) and inflammatory signaling pathways (e.g. TNF and NFκB), were downregulated by BYL719 (Figure 6-Figure Supplement 1A and B). Therefore, we hypothesized that, in addition to preventing the activation of osteogenic and chondrogenic differentiation pathways in progenitor cells, part of the mechanism by which BYL719 prevents HO in vivo may be due to the modulation of inflammatory responses.

### PI3K***α*** inhibition reduces the hyperinflammatory response in heterotopic ossification

To confirm whether BYL719 can modulate excessive inflammation at the injured sites, we examined its impact on monocytes, macrophages and mast cells *in vivo*. These cell populations are known to increase in number two to four days after injury (Convente et al., 2018; Sorkin et al., 2020), which coincides with the effects observed following early administration of BYL719. Hindlimb muscle samples were collected at two, four, nine, sixteen and twenty-three days post-induction of HO with Cre viruses and cardiotoxin in *ACVR1^Q207Dfl/fl^*mice (Figure 7A). The group of treated animals received BYL719 intermittent treatment starting one day after injury. After sixteen days, structures of mineralized bone became detectable in vehicle-treated mice (Figure 7-Figure Supplement 1A and B). The number of F4/80-positive monocytes/macrophages was high two days after injury and remained elevated in the HO lesions throughout the ossification process (Figure 7B and C). BYL719-treated animals showed an almost 50% reduction in the number of F4/80 positive cells at day two post-injury onwards. Consistent with previous studies, we observed an increased number of mast cells at injury sites. Their numbers peaked at four days and only dropped twenty-three days after injury in vehicle-treated mice (Figure 7D and E). In BYL719-treated mice, mast cells were increased up to day 9, but their number was completely normalized to preinjury levels already at day 16 (Figure 7D and E). This observation is consistent with the complete muscle regeneration observed in BYL719-treated mice compared to untreated mice, which are actively developing ossifications at that time (Figure 7-Figure Supplement 2). These results confirm a functionally relevant dual role of PI3Kα inhibition, by preventing the differentiation of osteochondrogenic progenitor cells and also by reducing the local inflammatory response.

**Figure 7.**
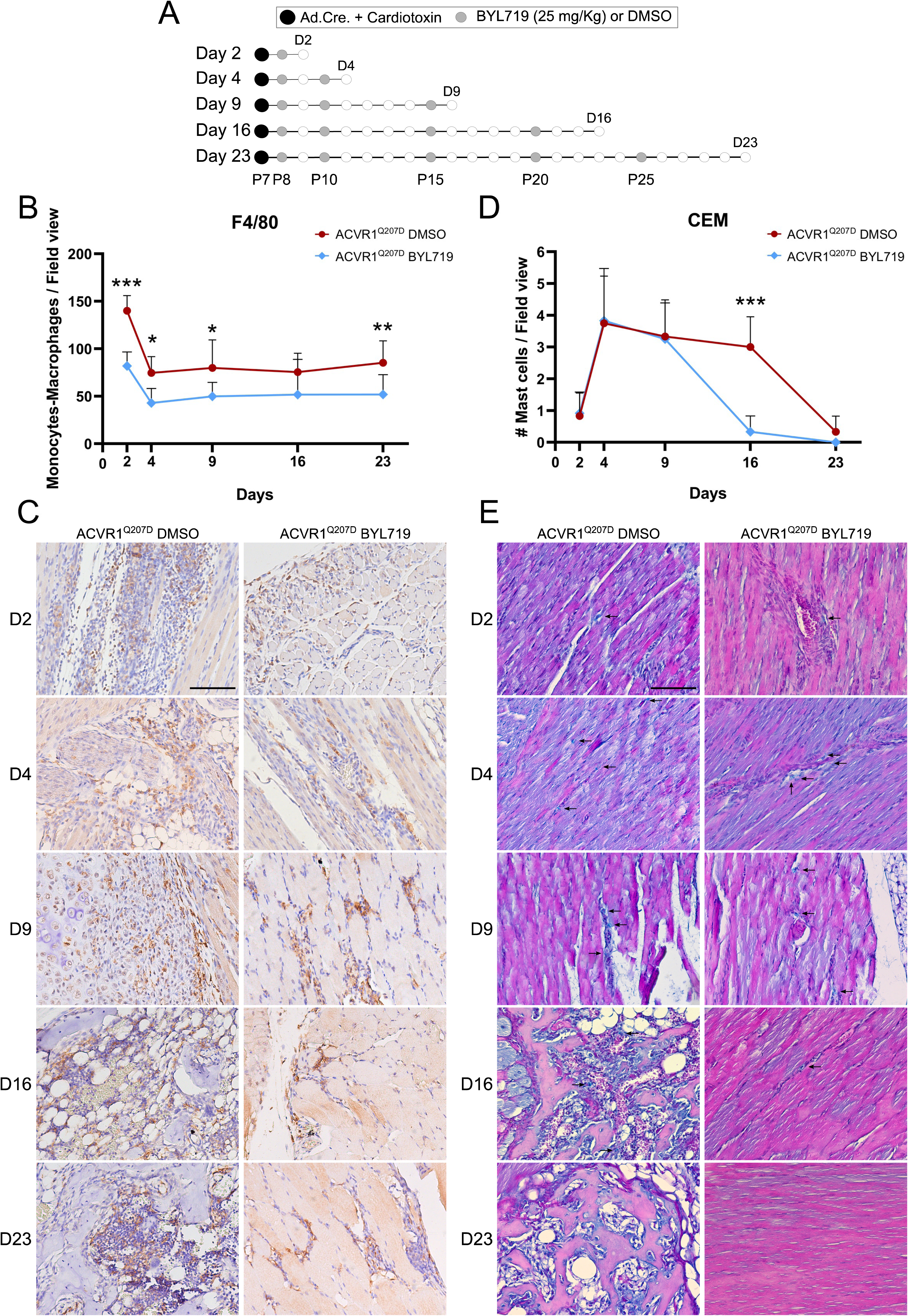
(A) Heterotopic ossification was induced at P7 through the injection of Adenovirus-Cre (Ad.Cre) and cardiotoxin in the mice hindlimb. Either DMSO (vehicle) or BYL719 (25 mg/kg) were injected following the scheme indicated with grey dots, starting at P8 but with a different duration P9 (2 days post-HO-induction), P11 (4 days post-HO-induction), P16 (9 days post-HO-induction), P23 (16 days post-HO-induction), and P30 (23 days post-HO-induction). (B) The quantification of F4/80-positive monocytes/macrophages per field view from immunohistochemistry staining, with 20x amplification as shown in the representative images depicted in Figure 8C. Five images were randomly acquired per mice, from 2 mice per group, for a total of 10 quantified images per group. F4/80 positive cells were detected as cells with dark brown staining. Data are shown as mean ± SD. *P<0.05, **P<0.01, ***P<0.001, two-way ANOVA with Sidak’s multiple comparisons test, comparing both groups on each day (n=10). (C) Representative images from the quantified immunohistochemistry staining for F4/80 to detect monocytes/macrophages. Representative images are shown for each final time-point at day 2, 4, 9, 16, and 23 post-HO-induction. All images were acquired at 20x (scale bar 100 µm). (D) Quantification of CEM-positive mast cells per field view from CEM staining, with 20x amplification as shown in the representative images depicted in Figure 8E. Three images were obtained per mice, from 4 mice per group, for a total of 12 quantified images per group. Mast cells were detected as cells with bright blue staining. Data are shown as mean ± SD. ***P<0.001, two-way ANOVA with Sidak’s multiple comparisons test, comparing both groups in each day (n=12). (E) C.E.M. staining was performed to detect mast cells, highlighted with black arrows. Representative images are shown for each final time-point at day 2, 4, 9, 16, and 23 post-HO-induction. All the images were obtained at 20x magnification, with a representative scale bar at 100 µm.

### PI3K***α*** inhibition reduces proliferation, migration and inflammatory cytokine expression in monocytes, macrophages and mast cells

Next, we used several models of disease-relevant immune cells to study the effect of pharmacological PI3Kα inhibition. Given the technical difficulties to transduce immune cells with lentiviral particles carrying ACVR1 R206H, we decided to partially recapitulate ACVR1 R206H activation with recombinant BMP6 and to test the effect of BYL719 in these conditions. Activation of ACVR1 signaling with BMP6 did not significantly modify the proliferation rate of any of the cell lines tested, that is, monocytes (THP1), macrophages (RAW264.7) and mast cells (HMC1) (Figure 8A, B and C). However, incubation with BYL719 in vitro strongly reduced their growth rate at 2µM and almost completely arrested their proliferation at 10µM. It is well known that monocytes are actively recruited from circulation to the ossification sites early after injury (Sorkin et al., 2020). We also examined the effects of BYL719 in proliferation in additional cell types involved in HO, such as myoblast and mesenchymal progenitors. BYL719 reduced the proliferation of myoblast and mesenchymal progenitors *in vitro* (Figure 8, Figure Supplement 1 A and B). However, the reduction in the proliferation did not reach the extent observed in monocytes or macrophages. We expanded our analysis by interrogating chemotactic migration of monocytes in transwell assays. Whereas monocytes actively migrated towards FBS as a chemotactic agent, the incubation of monocytes with BYL719 deeply diminished their migratory ability (Figure 8D). These results suggest that BYL719 is able to diminish monocyte recruitment, while reducing also the expansion of inflammatory cells at injury sites.

**Figure 8.**
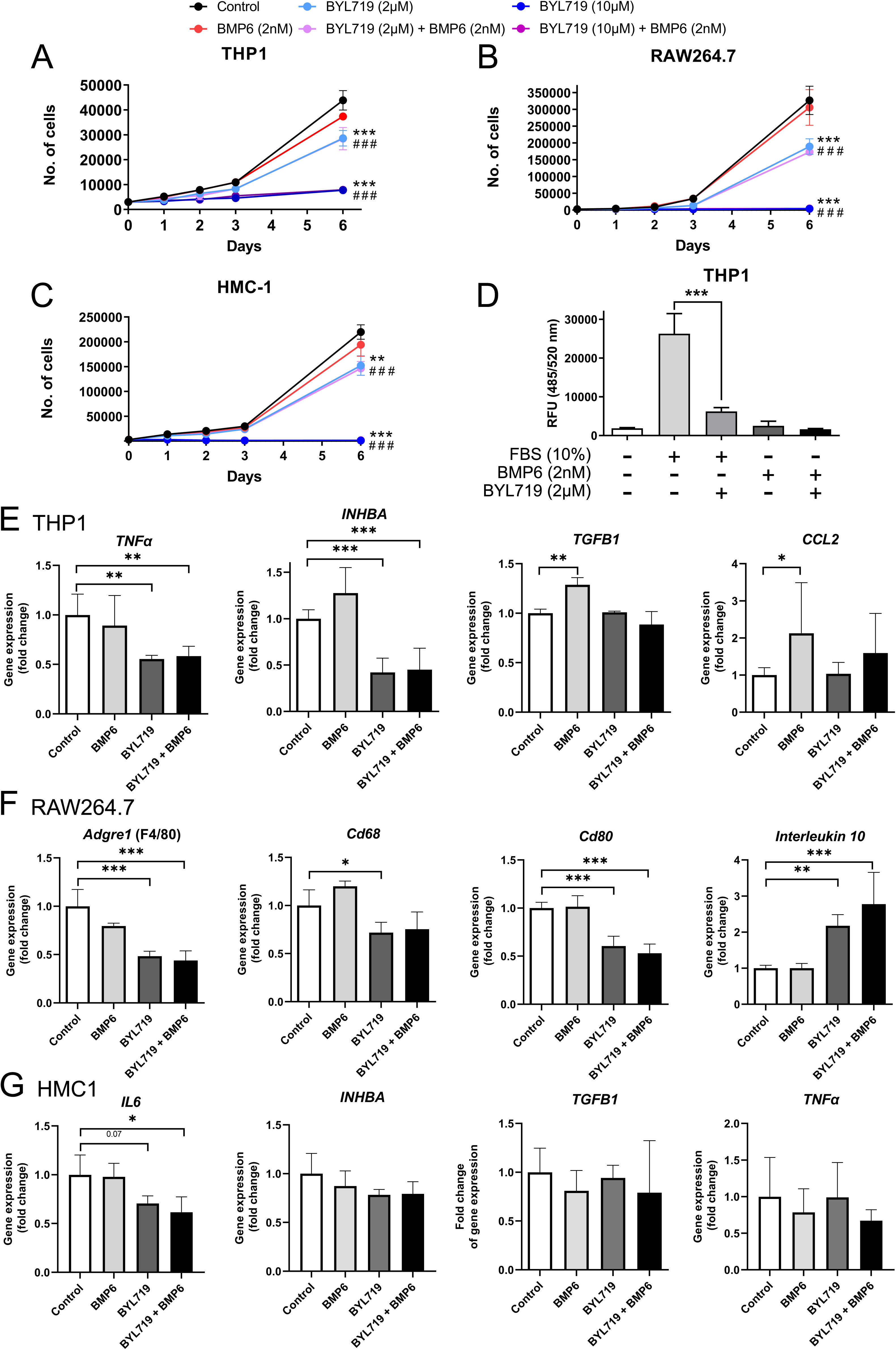
(A, B, C) Proliferation assays of THP1 (A), RAW264.7 (B), and HMC-1 (C) cells. Cells were cultured for 6 days and in control conditions, with BMP6 (2nM) and/or BYL719 (2 or 10µM). Areas under the proliferation curves were compared by one-way ANOVA with Dunnett’s multiple comparisons test against the control. (*) refers to significant differences between control and single treatments (BMP6 or BYL719). (#) refers to significant differences between control and combined treatments. Data shown as mean ± SD (n = 4 per group). ** or # # P<0.01, *** or # # # P<0.001. (D) Migration assay of THP1 cells with control conditions, or with FBS or BMP6 as chemotactic agents with or without BYL719 treatment. Data are shown as mean ± SD (n = 4 per group). ***P<0.001, two-way ANOVA with Tukey’s multiple comparisons test. (E, F, G) Gene expression assays of THP1 (E), RAW264.7 (F) and HMC-1 (G) cells. Cells were treated for 48 hours with BYL719 (2µM) and/or BMP6 (2nM). Data are shown as mean ± SD (n = 4 per group). *P<0.05, **P<0.01, ***P<0.001, one-way ANOVA with Dunnett’s multiple comparisons test, significance shown between control group and other groups.

Upon tissue injury, a multitude of pro-inflammatory and anti-inflammatory cytokines are released at the site of injury. This triggers the recruitment and infiltration of various cell types, including immune cells and osteochondrogenic progenitors. We examined the expression of distinct cytokines upon ACVR1 stimulation with BMP6 in the presence of BYL719 in monocytic, macrophagic and mast cell lines. In monocytes, BYL719 inhibited the expression of the pro-inflammatory cytokine *TNF-*α and the expression of activin A (*INHBA*), whereas the expression of transforming growth factor *TGF-*β*1* was increased by BMP6 and not modified by the addition BYL719 (Figure 8E). *CCL2* (MCP-1), a chemokine inducing the recruitment of monocytes and macrophages into inflamed tissues as well as the M2 polarization of macrophages (Sierra-Filardi et al., 2014), was significantly induced by BMP6 and partially reduced by BYL719 (Figure 8E). In macrophages, BYL719 reduced the expression of pan-macrophage markers *Adgre1* (F4/80), *Cd68* and the M1 macrophage marker *Cd80* (Figure 8F). Conversely, BYL719 increased the expression of the M2 macrophage marker *Il10*, regardless of the presence or absence of BMP6. In the human mast cell line HMC1, BYL719 inhibited the expression of the pro-inflammatory cytokine *IL6*, while the expression of *TGF*β*1*, *TNF*α, and activin A (*INHBA*) remained unchanged upon the addition of either BMP6 or BYL719 (Figure 8G). We also analyzed the effects of BYL719 on myogenic differentiation *in vitro*. Addition of 2 or 10µM BYL719 was able to abolish the expression of the myogenic markers *MyoD1* and myosin heavy-chain 1 (*Myh1*) while barely altering the expression of the myofibroblast marker α-SMA (*Acta2*) (Figure 8-Figure Supplement 1C). Similarly, BYL719 (at 2 or 10µM) was able to block formation of myotubes after 7 days of myogenic differentiation *in vitro* (Figure 8-Figure Supplement 1D). Altogether, these data suggest that PI3Kα inhibition might reduce the expression of certain pro-inflammatory cytokines. Moreover, even though these results should be further validated in cells carrying mutated ACVR1, the reduction on activin A expression by BYL719 observed in inflammatory cells and mesenchymal progenitors could be relevant for HO in FOP.

## Discussion

We previously showed that pharmacological PI3Kα inhibition using BYL719 has the potential to suppress HO by increasing SMAD1/5 degradation, reducing transcriptional responsiveness to BMPs, and blocking non-canonical responses such as the activation of AKT/mTOR (Valer et al., 2019a). In this manuscript, we confirmed these results through both pharmacological and genetic approaches, and further expanded the insights into the molecular and cellular mechanisms responsible for the therapeutic effect of PI3K inhibition. Here, we optimized the therapeutic window for BYL719 administration and found that the delayed administration of BYL719 effectively prevents HO. Moreover, *in vivo* simultaneous genetic inhibition of *p110*α and the activation of mutated *ACVR1* at injury sites partially prevents HO, and we demonstrate that inhibition of PI3Kα inhibits osteochondroprogenitor specification. We determined that BYL719 does not inhibit recombinant ACVR1^R206H^ kinase activity *in vitro* and does not behave as an ATP competitor inhibitor for none of the TGF-β receptor kinases. Finally, the administration of BYL719 prevented an exacerbated inflammatory response *in vivo*, possibly due to the effects observed on immune cell populations.

While FAPs play a key role in bone formation in different types of HO, inflammation is also closely associated with all types of HO (Barruet et al., 2018; de Ruiter et al., 2021; Matsuo et al., 2019). For instance, the depletion of mast cells and macrophages has been shown to reduce bone volume in FOP mouse models (Convente et al., 2018. Moreover, Activin A, TGF-β and other cytokines secreted by FAPs, monocytes, macrophages and mast cells are essential for FOP and non-genetic HO (Alessi Wolken et al., 2017; Hatsell et al., 2015; Lees-Shepard et al., 2018; Patel et al., 2022; Sorkin et al., 2020; Upadhyay et al., 2017). In addition, ACVR1 antibodies can activate ACVR1^R206H^ even in the absence of Activin A, but muscular trauma is still required to induce HO (Aykul et al., 2022; Lees-Shepard et al., 2022), indicating the involvement of additional immune factors. Our study finds that BYL719, a PI3Kα inhibitor, reduces the expression of certain pro-inflammatory cytokines, effectively blocks the proliferation of monocytes, macrophages and mast cells and reduces the migratory potential of monocytes. This likely contributes to a decreased number of monocytes and macrophages at injury sites and throughout the in vivo ossification process. It is known that excessive pro-inflammatory cytokine expression, including activin A, by monocytes and macrophages is observed in all types of HO and is induced by ACVR1^R206H^ (Matsuo et al., 2021). Non-canonical signaling pathways (mTOR, p38, TAK1, NF-κB) are also affected by ACVR1^R206H^ in immune cell types (Barruet et al., 2018; Hwang et al., 2022). M1-type macrophages initiate acute inflammatory responses but later transition to M2-type macrophages associated with anti-inflammatory and reparative functions. Therefore, modulating macrophage phenotype towards regeneration may dampen HO (Sorkin et al., 2020). BYL719 reduces monocyte/macrophage numbers, migration, and pro-inflammatory cytokine expression, potentially altering their polarization. It also reduces activin A expression and promotes a shift towards the M2 phenotype. BYL719 can help to reduce the hyper-inflammatory state in HO, inhibit FAP expansion, and favor myogenic regeneration of muscle tissue (Stanley et al., 2022).

Our observation that the delayed administration of BYL719 is still effective has pathophysiological and therapeutic derivatives. Muscle injury in mice induces sequential changes in HO progression: early immune cell infiltration (days 1-3), a fibroproliferative and late inflammatory phase, with an expansion of FAPs and a spectrum of monocytes, macrophages and mast cells as highly secretory cells (days 3-7), followed by chondrogenesis (day 7-14) and osteogenesis (days 14-23) (Hwang et al., 2022). Starting BYL719 administration 3 days post-injury is still fully therapeutically effective and even partially at 7 days post-injury. In addition, whereas untreated mice developed cartilage and bony lesions by days 9 and 16 post-injury, respectively, BYL719-treated animals fully regenerated muscle by day 16 post-injury. Altogether, this evidence suggests that the major effects of BYL719 occur between 3 and 16 days post-injury. We found that BYL719 completely blocks chondrogenesis in cultured hMSCs. In addition, in the histological analyses of these BYL719-treated mice that do not develop HO, we cannot find any sign of cartilage formation on either the 9^th^, 16^th^ or 23^rd^ days post-injury, which confirms that BYL719 is also able to block chondrogenesis in mice *in vivo*. This suggests that intervention with BYL719 over a temporal window after a HO flare might be sufficient to inhibit endochondral ossification. In addition, although surgical resection is not recommended in FOP patients and has a risk of recurrence in resected non-genetic HO, it could be envisaged using as preventive therapy in subjects undergoing surgeries to remove ectopic bone.

Treatment options for HO and FOP are currently limited, primarily consisting of anti-inflammatories such as corticosteroids and NSAIDs. Promising pharmacological treatments have progressed to II/III clinical trials. Palovarotene, a retinoic acid receptor-γ agonist, has been approved in Canada and U.S.A. for patients over 10 years old (NCT03312634). However, a previous clinical trial with palovarotene was paused for children due to concerns about premature growth plate closure, which is debated in FOP mice (Chakkalakal et al., 2016; Rosen et al., 2018)(Chakkalakal et al., 2016; Rosen et al., 2018). Antibodies targeting activin A (Garetosmab) showed effectiveness, but serious adverse effects led to a temporary hold on the trial (NCT03188666). The outcome of a clinical trial with Rapamycin has not been publicly disclosed and a case report indicated limited benefits in classical FOP patients (UMIN000028429) (Kaplan et al., 2018). BYL719 (Alpelisib/Piqray®) was recently licensed in combination with hormone therapy for a PIK3CA-mutated breast cancer (Cardoso et al., 2020). BYL719 has been shown to be also clinically effective in adult patients with PIK3CA-related overgrowth syndrome (PROS) and children under 1 year of age (Madsen and Semple, 2022; Morin et al., 2022). In both scenarios, BYL719 had no major adverse effect when patients were treated for more than one year (in adults at a dose of 300mg daily for oncological or PROS purposes and at a dose of 25 mg daily in infants with PROS) (Morin et al., 2022). Thus, implementing BYL719 for treatment of HO, at least during a restricted temporal window (i.e. surgery to remove ectopic bone in HO, or during flare-ups in FOP individuals), might be a valid therapeutic option for FOP patients.

## Materials and Methods

### Murine bone marrow mesenchymal stem cells isolation

Murine bone marrow-derived mesenchymal stem cells (BM-MSCs) were isolated from 6-8 weeks-old C57BL6/J mice for the floxed p110α allele (Beatriz Gámez et al., 2016) as previously described (Soleimani and Nadri, 2009). We also isolated BM-MSCs from UBC-CRE-ERT2/ACVR1^R206H^ *^fl/wt^* (a gift from Dr. Dan Perrien and IFOPA). This conditional ACVR1^R206H^ construct was knocked-in to the endogenous *Acvr1* gene immediately following intron 4. After 4=OH tamoxifen addition, CRE activity excises murine *Acvr1* exons 5-10 and induce expression of the corresponding exons of human ACVR1^R206H^ and an eGFP marker. In both cases, mice were euthanized and femurs were dissected and stored in DMEM with 100 U/ml penicillin/streptomycin (P/S). Soft tissues were cleaned and femur ends were cut under sterile conditions. Bone marrow was flushed with media using a 27-gauge needle. The resulting cell suspension was filtered through a 70-μm cell strainer and seeded in a 100-mm cell culture dish. Non-adherent cells were discarded after 3 hours. Media was slowly replaced every 12 hours for up to 72 hours. Then, media was replaced every 2 days until the culture reached 70% confluence. Then, cells were lifted by incubation with 0.25% trypsin/0.02% EDTA for 5 minutes at room temperature. Lifted cells were cultured and expanded. MSCs from UBC-CRE-ERT2/ACVR1^R206H^ *^fl/wt^* mice were treated with 4-hydroxytamoxifen to induce Cre recombination.

### Production of human mesenchymal stem cells (hMSCs) expressing ACVR1

For chondrogenic differentiation and RNA sequencing, bone marrow-derived human MSCs were transduced by lentiviral delivery encoding ACVR1^WT^ or ACVR1^R206H^ (Van Dinther et al., 2010). For NanoBRET target engagement kinase assays, were transduced with ACVR1^WT^-nanoluc or ACVR1^R206H^-nanoluc. The nanoluc fusion inserts (Promega, NV2341/NV2381) were subcloned into a PLV-CMV-IRES vector. First, an eGFP was amplified with PCR containing PstI, SalI and XbaI restriction sites and subcloned in the PLV using PstI and XbaI. Next, the ACVR1-nanoluc inserts were subcloned in the PLV by restriction of PstI and SalI, exchanging the eGFP. Subsequently, lentiviral particles were produced using HEK293t cells and cells were transduced.

### Cell culture

Murine BM-MSCs were cultured in DMEM supplemented with 10% FBS, 2 mM glutamine, 1 mM sodium pyruvate, and 100 U/ml P/S and incubated at 37°C with 5% CO_2_. The human monocyte cell line THP1 was cultured in RPMI-1640 supplemented with 10% FBS, 2 mM glutamine, 1 mM sodium pyruvate, 100 U/ml P/S, 0.1mM NEAA, 10mM HEPES and 50µM 2-mercaptoethanol. Mouse macrophage cell line RAW264.7 was cultured in DMEM supplemented with 10% FBS, 2 mM glutamine, 1 mM sodium pyruvate and 100 U/ml P/S. The human mast cell line HMC1 was cultured in IMDM supplemented with 10% FBS, 2 mM glutamine, 1 mM sodium pyruvate and 100 U/ml P/S. The hMSCs were cultured in αMEM supplemented with 10% FBS, 0.1 mM ascorbic acid, 1 ng/ml human bFGF and 100 U/ml P/S (Growth Medium; GM). C2C12 cells were cultured in DMEM supplemented with 20% FBS, 2 mM glutamine, 1 mM sodium pyruvate, and 100 U/ml P/S and incubated at 37°C with 5% CO_2_. For C2C12 differentiation, cells were cultured for seven days in the same media with 5% horse serum.

### Proliferation and chemotactic migration assays

For proliferation assays, cellswere seeded in 12-well culture plates in correspondent culture medium with 2 nM BMP6 and/or 2 μM or 10 μM BYL719 (Chemietek). Photos of several fields of each well were taken at 24, 48, 72 and 144 hours using a Leica DM-IRB inverted microscope. Cell counting was performed through ImageJ software.

The chemotactic migration assay of THP1 was performed through the InnoCyte Monocyte Migration Assay (Merck Millipore). Cells were serum-starved for 3h prior to the experiment and, when needed, treated with 2 μM BYL719 for 30 minutes before the experiment started. Then, 10^5^ cells/100 μL were added to each upper chamber. Lower chambers were charged with 10% FBS-supplemented RPMI-1640 (positive control), FBS-depleted RPMI-1640 (negative control) or 2 nM BMP6 (R&D), respectively. Cells migrated for 2h and were then stained with calcein-AM and pipetted into a 96-well conical bottom plate (Sigma-Aldrich). A plate reader was used to measure florescence from the top of the plate at an excitation wavelength of 485 nm and an emission wavelength of 520 nm. Unless otherwise stated, BYL719 at 2 or 10 μM (Chemietek) and 2 nM BMP6 (R&D) was used for 6 days (proliferation) or 4 hours (migration) in HMC1, THP1 and RAW264.7 cells.

### Retroviral transduction and gene expression analysis

BM-MSCs were infected with mock virus (pMSCV) or with viruses expressing wild type *Acvr1* (WT), *Acvr1*^R206H^ (RH) and/or pMSCV-Cre. Plasmids with WT and RH *Acvr1* forms were kindly provided by Dr. Petra Seemann and were subcloned into pMSCV vector. *Acvr1* and p110α expression levels were analyzed by qRT-PCR. Cells were treated with BYL719 (2µM) and/or activin A (2nM) for 48 hours in complete media without FBS. THP1, RAW264.7, and HMC-1 cell lineswere treated in the media indicated for each cell line with BYL719 (2µM) and/or BMP6 (2nM) for 48 hours.

Total RNA for RT-PCR was extracted from all cellular models using TRIsure reagent. At least 2 μg of purified RNA was reverse-transcribed using the High-Capacity cDNA Reverse Transcription Kit (Applied Biosystems). Quantitative PCRs were carried out on ABI Prism 7900 HT Fast Real-Time PCR System with Taqman 5′-nuclease probe method and SensiFAST Probe Hi-ROX Mix. All transcripts were normalized using *Tbp* as an endogenous control.

### NanoBRET target engagement assays

COS-1 cells were cultured in DMEM (Gibco, 11965092) supplemented with 10% FBS and 100 U/ml P/S. These cells were transfected in 70% confluent 6-wells TC-treated plates with 2 µg TGFβ receptor-Nanoluciferase constructs (constructs were provided by Promega and Promega R&D) using a 1 µg DNA: 2 µl PEI (1 mg/ml) ratio. The transfected cells were reseeded in white 384-wells TC-treated assay plates one day before the nanoBRET readout in DMEM supplemented with 1% FBS in a quantity of 15*10^3 cells in 40 µl per well. Two hours before measurement, the dedicated tracer (Promega) and the test compounds were incubated to the cells at 37°C. Immediately after intracellular TE Nano-Glo Substrate/Inhibitor (Promega, N2160) addition, the wells were measured by 450-80BP and 620-10BP filters using the ClarioSTAR (BMG Labtech). Subsequent nanoBRET ratios were measured by the formula: acceptor emission (620-10 nm)/donor emission (450-80 nm) * 1000 (milliBRET units, mBU).

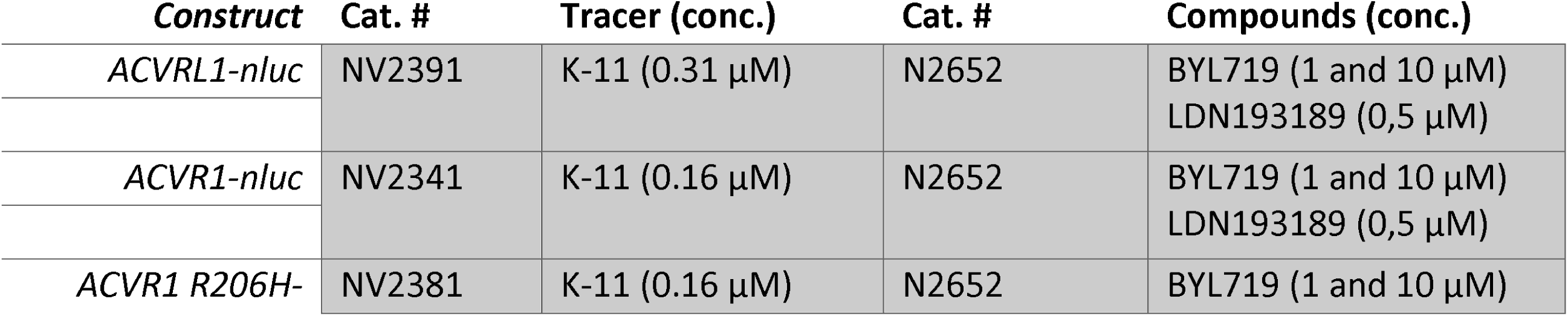
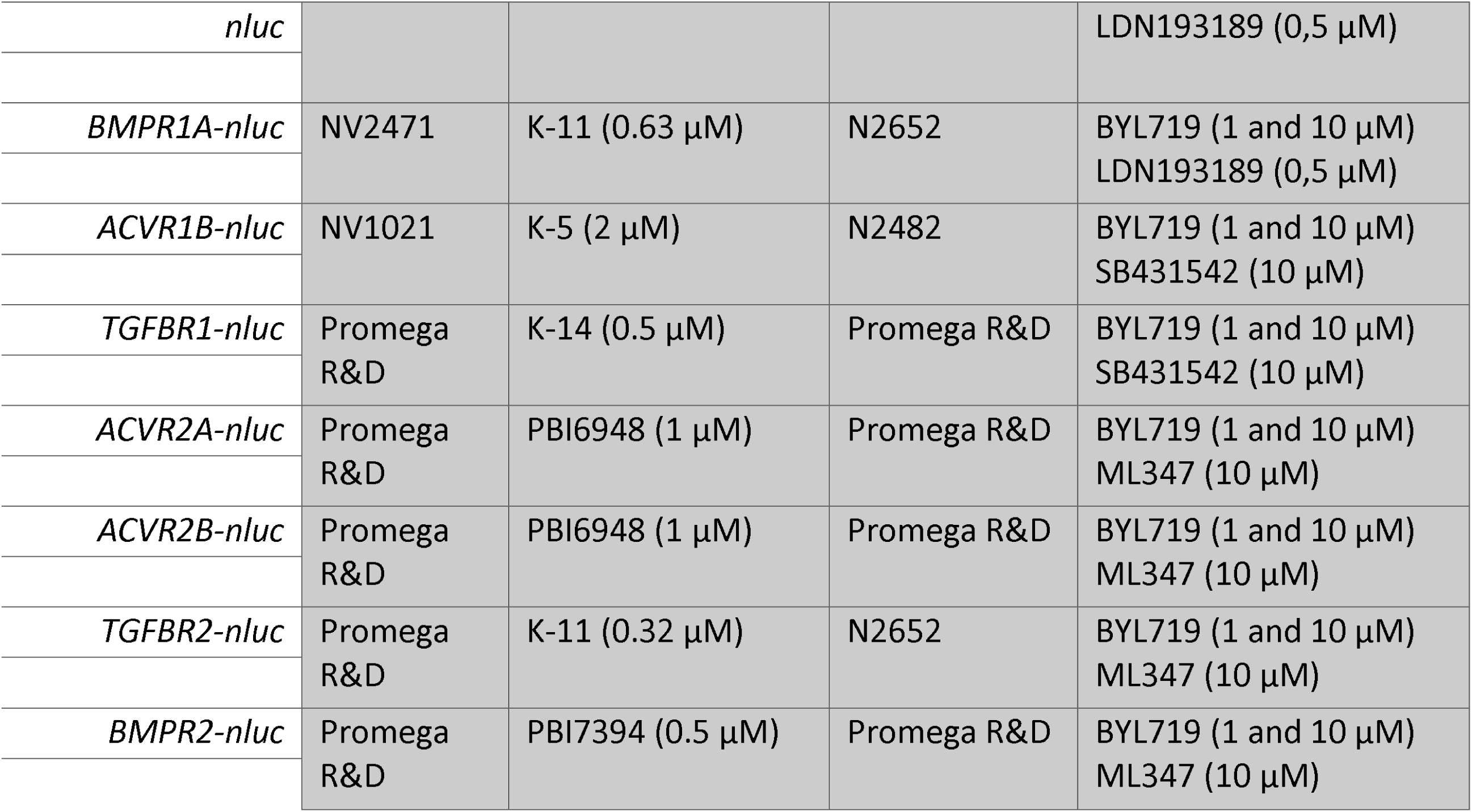

### Chondrogenic progenitor differentiation

MSCs were cultured in αMEM supplemented with 10% FBS, 1 ng/ml human bFGF and 100 U/ml P/S (Growth Medium; GM). Chondrogenic differentiation medium (DM) consisted of DMEM/F12 supplemented with 1% Insulin-Transferrin-Selenium, 170 µM Ascorbic Acid, 0.1 µM Dexamethasone, 350 µM L-proline, 1 mM Sodium Pyruvate, 0.15% glucose, 1% FBS and 100 U/ml P/S. During differentiation, the medium was refreshed twice a week for the duration of the experiment. For the Alcian Blue experiments, micromass cultures were made by carefully seeding 3×10^5^ cells per 10 µl droplets in a 24-wells plate and incubated for 2 hours at 37°C prior to GM addition. Chondrogenic progenitor differentiation was started 1 day after micromass seeding by the addition of DM. Dependent on the condition, we supplemented the DM with TGF-β1 (10 ng/ml) or Activin A (100 ng/ml), and treated with BYL719 (10 µM) or DMSO vehicle. For the ALP staining, the hMSC-ACVR1 lines were seeded at 2×10^4^ cells per well in 48-wells plates and differentiated for 7 and 11 days.

### Alkaline phosphatase (ALP) staining, ALP activity assay and Alcian Blue staining

For the ALP staining, the cells were washed with PBS, fixated using 3.7% formalin for 5 minutes at RT, washed twice with PBS and stained in ALP solution containing 2 mg Naphthol AS-MX, 6 mg Fast Blue, 5 ml 0.2M Tris (pH 8.9), 100 µl MgSO_4_ and dH_2_O up to 10 ml. Images were acquired using the Leica DMi8 with a 10x magnification. To measure ALP activity, ALP lysates were obtained by washing twice with PBS, freezing at -80°C for 1 hour, and lysing on ice for 1 hour using 100 µl ALP buffer (100 µM MgCl2, 10 µM ZnCl2, 10 mM Glycine (pH 10.5)) plus 0.1% Triton X-100 per well. ALP activity was quantified by adding 20 µl lysate and 80 µl 6 mM PNPP in ALP buffer to a clear 96-well plate incubated at RT until the samples turned yellow. Next, the samples were measured at an absorbance of 405 nm.

The micromass pellets were stained with Alcian Blue as described before (Sánchez-Duffhues et al., 2019). Images were acquired using the Leica DMi8 with 5x magnification. Densitometry analysis of the whole well (from total plate scan) was performed using ImageJ (v1.53t).

### Bulk RNA sequencing and analyses

The hMSCs-ACVR1^WT^ and hMSCs-ACVR1^R206H^ were seeded at 100% confluency and serum starved overnight with DMEM without supplements. Afterwards, the cells were pre-treated with or without 1 µM of BYL719 in starvation medium for 30 minutes and stimulated with or without 50 ng/ml Activin A or 50 ng/ml BMP6 for 1 hour. RNA was isolated using the ReliaPrep RNA Miniprep Systems (Promega) and sequenced using the Illumina NovaSeq 6000 (Illumina) platform. The reads were mapped using the Hisat2 (v2.0.5) package and the total and normalized counts were measured using featureCounts (v1.5.0-p3). Differential expression analysis was performed using the DESeq2 R package (v1.20.0), in which the p-values were adjusted following the Benjamini and Hochberg approach. Genes with an adjusted p-value < 0,05 were considered differentially expressed. Subsequent Gene Ontology analysis was performed using the clusterProfiler R package and the GO terms were considered significant if the p-values < 0,05. Gene Set Enrichment Analysis was performed using the GSEA analysis tool (gsea-msigdb.org). Heatmaps were generated using heatmap2 function within the R gplots package in Galaxy (usegalaxy.eu). The datasets are publicly available (see GSE237512).

### Western blot

Cells were lysed using RIPA lysis buffer and quantified using the Pierce BCA Protein assay (Thermo) following standard protocols. The protein samples were loaded using β-ME containing sample buffer and run using SDS-PAGE and transferred to PVDF membranes. Primary antibodies against SMAD1 (Cell Signaling, 6944), p-SMAD1/5 (Cell Signaling, 9516), and vinculin (Sigma, V9131) were used at 1:1000 dilution in 5% BSA in TBST. Binding was detected with HRP-conjugated secondary antibodies and visualized by Brightfield ECL (Thomas Scientific) on the ChemiDoc (BioRad).

### In vitro recombinant kinase activity assay

Fifty ng of recombinant ACVR1^R206H^ (ab167922, Abcam) was diluted in 30 µl of 25mM Tris/HCl (pH7.5), 10mM MgCl_2_, 10mM MnCl_2_ and 2mM DTT. The reaction was started by the addition of 5µM ATP-Mg, 0.05 µCi/µl ATP [γ-^32^P] and 0.5mg/ml of dephosphorylated casein (Sigma) and performed for 30 min at 30°C. The reaction was stopped and samples were processed by SDS-PAGE. After drying the gel, phosphorylation was determined by autoradiography.

### Heterotopic ossification mouse model

To study heterotopic ossification *in vivo*, we used the Cre-inducible constitutively active ACVR1^Q207D^ (CAG-Z-EGFP-caALK2) mouse model as previously described (Fukuda et al., 2006; Shimono et al., 2011; Yu et al., 2008). Mice ACVR1Q207D^fl/fl^ p110α^fl/fl^were obtained by crossing the detailed homozygous ACVR1Q207D^fl/fl^ with the p110α^fl/fl^homozygous mutant mice carrying loxP sites flanking exons 18 and 19 of the p110α alleles (Beatriz Gámez et al., 2016). Heterozygous mice were crossed until ACVR1Q207D^fl/fl^ p110α^fl/fl^was obtained.

To induce heterotopic ossification in P7 ACVR1^Q207D^ mice, 1×10^8^ pfu of adenovirus-Cre (Ad-CMV-Cre, Viral Vector Production Unit, UAB) and 0.3 μg of cardiotoxin in 10 μl 0.9% NaCl volume were injected into the left hindlimb. Control groups had the same procedure without Ad-Cre in the injection. On P8, mice started receiving either placebo (intraperitoneal administration (i.p.) of DMSO) or BYL719 (i.p. of 25 mg/kg), both diluted in 0.5% carboxymethylcellulose sodium. Mice were housed under controlled conditions (12-hour light/12-hour dark cycle, 21°C, 55% humidity) and fed *ad libitum* with water and a 14% protein diet (Teklad2014, Envigo). Mice were regularly weighted over the whole period. Heterotopic ossification induction was performed blinded to mouse and group identity. All procedures were approved by the Ethics Committee for Animal Experimentation of the Generalitat de Catalunya.

### Micro-computed tomography analysis

The caudal half of mice was collected and fixed in 4% paraformaldehyde (PFA) for 48 hours at 4°C. Samples were conserved in PBS and high resolution images were acquired using a computerized microtomography imaging system (Skyscan 1076, Bruker microCT), in accordance with the recommendations of the American Society of Bone and Mineral Research (ASBMR). Samples were scanned in air at 50 kV and 200 µA with an exposure time of 800 ms, using a 1 mm aluminum filter and an isotropic voxel size of 9 µm. Two dimensional images were acquired every 1° for 180° rotation and subsequently reconstructed, analyzed for bone parameters, and visualized by NRecon v1.6, CT-Analyser v1.13 and CTVox v3.3 programs (Bruker), respectively. For heterotopic ossification, manual VOIs comprising heterotopic ossifications were employed and a binary threshold was established at 25-255.

### Histology and immunoflouresecence

Whole legs were fixed in 4% PFA for 48 hours at 4°C, decalcified in 16% EDTA pH 7.4 for 6 weeks, and embedded in paraffin. Five µm sections were cut and stained with Fast Green/Safranin O, or Masson’s Trichrome. Images were obtained with brightfield Eclipse E800 (Nikon). To detect monocyte/macrophage cells, paraffined slides were immunostained with Anti-F4/80 antibody [SP115] (ab111101). Briefly, after deparaffination and rehydration, an antigen retrieval step with citrate buffer at pH6 in a decloaking chamber was applied to the slides. Slides were cooled down and rinsed with 1x TBS and endogenous peroxidases were blocked with 70% methanol, 28% distilled water, and 2% hydrogen peroxide for 5 minutes at room temperature. After a blocking step, F4/80 primary antibody incubation at concentration 1:100 overnight at 4°C was applied. Envision Dual Link conjugated with HRP was applied for 40 minutes at room temperature, and detection was performed with diaminobenzidine (DAB) for 3 minutes. Nuclear staining with Lilly’s hematoxylin was applied to the slides for 30 seconds. Images of stained slides were obtained with brightfield Eclipse E800 (Nikon). Images were randomly acquired with 20x magnification from the region of interest (injected muscle, with or without visible heterotopic ossification). Five images were obtained per mice, from 2 mice per group, for a total of 10 quantified images per group. F4/80 positive cells were detected as cells with dark brown staining. To detect mast cells, paraffined slides were stained with C.E.M. staining using the Eosinophil-Mast cell staining kit (ab150665, Abcam). The recommended staining procedure was applied to the deparaffinized sections. Images of stained slides were obtained with brightfield Eclipse E800 (Nikon). Images were randomly acquired with 20x magnification from the region of interest (injected muscle, with or without visible heterotopic ossification). Three images were obtained per mice, from 4 mice per group, for a total of 12 quantified images per group. Mast cells were detected as cells with bright blue staining.

For immunofluorescence assays, five µm sections of each time point and condition were deparaffinased, processed with antigen retrieval and permeabilization steps, and stained with wheat germ agglutinin (1:250 dilution, ThermoFisher Scientific), followed by incubation using PDGFRA antibody (1:250 dilution Cell Signalling #3174), Alexa555 secondary antibody, and DAPI staining. Images were randomly acquired with 20x magnification from the region of interest with a Carl Zeiss Axio Imager M2 Apotome microscope. Five images were obtained per mice, from 4 mice per group, for a total of 20 quantified images per group.

### Statistical analysis

Unless stated otherwise in each figure legend, the results were expressed as mean ± SD. The median was shown in heterotopic ossification datasets where non-parametric tests were applied. Each figure legend has detailed information explaining the statistical test used to compare between groups, the compared groups, the explanation of the symbols used to show significance, usually *P<0.05, **P<0.01, ***P<0.001, and the sample size of the experiment. Heterotopic ossification, evaluation, and quantification was performed blinded to mouse and group identity. Microscopy and histology representative images were selected from the total quantified images. Microscopy and histology quantified fields were randomly acquired from the plate or the region of interest. Statistical tests were performed on GraphPad Prism 9.5.

## Acknowledgments

We thank E. Adanero, E. Castaño for technical assistance. We acknowledge the support from Promega R&D. Alexandre Deber is a recipient of a F.P.I. fellowship from the Ministry of Science and Innovation (MCIN/AEI/ 10.13039/501100011033). Carolina Pimenta-Lopes is the recipient of a F.P.U. fellowship from the Spanish Ministry of Education. This research was supported by grants PID2022-141212OA-I00, PDC2021-121776-I00 and PID2020-117278GB-I00 from MCIN/AEI/10.13039/501100011033, co-funded by FEDER ‘Una manera de hacer Europa’ and ‘NextGenerationEU’/PRTR; a grant 202038-30 from La Marató de TV3 and grants from IFOPA (ACT for FOP) and FOP Italia. MW and MJG are sponsored by the Netherlands Cardiovascular Research Initiative (the Dutch Heart Foundation, Dutch Federation of University Medical Centers, the Netherlands Organization for Health Research and Development, and the Royal Netherlands Academy of Sciences), PHAEDRA-IMPACT (DCVA) and DOLPHIN-GENESIS (CVON). GSD is also sponsored by the Spanish Ministry of Science through the Ramón y Cajal grant RYC2021-030866-I and the BHF-DZHK-DHF, 2022/23 award PROMETHEUS.

## Author contributions

J.A.V., A.D., M.W., M.J.G., J.L.R., G.S-D. and F.V. designed the study, and analyzed and interpreted the results. J.A.V, A.D., M.W. and C.P-L. performed the experiments. J.A.V., A.D., G.S-D. and F.V. wrote the manuscript. All authors contributed to the manuscript. G.S-D. and F.V. supervised the project.

## Supplementary Figure Legends

**Figure 1-Figure Supplement 1.** Mice body weight from P8 (one day post-HO-induction) to P30. Data are shown as mean ± SEM. Two-way ANOVA with Sidak’s multiple comparisons test.

**Figure 2-Figure Supplement 1.** (A) Mice body weight from P8 (one day post-HO-induction) to P30 (final time-point). Data are shown as mean ± SEM. Two-way ANOVA with Sidak’s multiple comparisons test. (B) Close-up 3D microtomography images of the injected hindlimbs of mice for each experimental group. White arrows show heterotopic ossification. (C) Two-dimensional representation of bone volume (BV) and bone volume/tissue volume (BV/TV) within heterotopic ossifications. Data shown are of each individual mouse with observed heterotopic ossification. Each symbol corresponds to an individual mouse. Colours indicate groups, as detailed in the legend.

**Figure 2-Figure Supplement 2.** Muscle sections of mice 23 days after injury stained with Masson’s Trichrome. Different genotypes and treatments specified in the figure. All images were acquired at 4x magnification, (scale bar 500 µm).

**Figure 3-Figure Supplement 1.** (A) Muscle sections of mice 4, 9, 16 and 23 days after induction of heterotopic ossification (HO). PDGFRA+ cells are indicated in red, wheat germ agglutinin staining is indicated in green and DAPI staining is indicated in blue. Different times and BYL719 treatment are specified in the figure. (B) Quantification of PDGFRA-positive FAPs obtained from five images obtained per mice, from 4 mice per group, for a total of 20 quantified images per group. (C) Quantification of myotube diameter estimated from wheat germ agglutinin staining obtained from five images per mice, from 4 mice per group, for a total of 20 quantified images per group.

**Figure 4-Figure Supplement 1.** Quantification of the TGF-β receptor-Nanoluciferase protein expression levels measured by the raw donor emission (excitation at 450-480 nm).

**Figure 5-Figure Supplement 1.** Human MSC-ACVR1^R206H^ are responsive upon Activin A (50 ng/ml) stimulation compared to hMSC-ACVR1^WT^ after overnight starvation. (A) Western blot analysis detecting C-terminal phosphorylated SMAD1/5, total SMAD1 and vinculin. (B) RT-qPCR analysis of the BMP target genes ID1 and ID3 (n=3 per group). Data are shown as the mean ± SD. * P<0.05, ** P<0.01, *** P<0.001, by two-way ANOVA with Tukeýs multiple comparison test.

**Figure 6-Figure Supplement 1.** GSEA of bulk RNA sequencing comparing hMSC-ACVR1^R206H^ cells treated with BYL719 (1 µM) with untreated controls, both stimulated with Activin A (50 ng/ml) (n=4 per group). (A) Significant down-regulated enrichment plots of the KEGG gene set TNF signaling pathway (HSA04668), NF-κB signaling (HSA04064) and GO gene set response to interleukin-6 (GO:0070741). (B) The top 40 most relevant genes from the enrichment plot of the TNF signaling pathway as derived from the leading edge subset.

**Figure 7-Figure Supplement 1.** (A) Quantification of heterotopic ossifications bone volume (mm^3^) of each experimental group. group for each final time-point at day D-2, 4, 9, 16, and 23 post-HO-induction. Colored symbols indicate the presence of heterotopic ossifications. Black symbols indicate the absence of heterotopic ossifications. Circles indicate DMSO-treated mice, diamonds indicate BYL719-treated mice, as detailed in the legend. Data shown are of each individual mouse with group median. **P<0.01, two-way ANOVA with Sidak’s multiple comparisons test. (B) Representative 3D microtomography images of the injected hindlimbs of mice from each experimental group. White arrows show heterotopic ossification.

**Figure 7-Figure Supplement 2.** (A) Representative images of injected hindlimbs stained with Fast green/safranin O (FGSO) staining. Representative images are shown for each final time-point at day 2, 4, 9, 16, and 23 post-HO-induction. All images were obtained at 4x (scale bar at 500 µm). (B) Representative images of injected hindlimbs stained with Masson’s trichrome. Representative images are shown for each final time-point at day 2, 4, 9, 16, and 23 post-HO-induction. All images were acquired at 4x magnification (scale bar at 500 µm).

**Figure 8-Figure Supplement 1.** (A and B) Proliferation assays of murine MSCs (A), and C2C12 cells (B). Cells were cultured for 3 days and in control conditions, or with BYL719 (2 or 10µM). Areas under the proliferation curves were compared by one-way ANOVA with Dunnett’s multiple comparisons test against the control. (C) RT-qPCR analysis of the myogenic (*MyoD1* and *Myh1*) or myofibroblast (*Acta2*) markers after culturing C2C12 cells in myogenic differentiation media for 7 days in control conditions, or with BYL719 (2 or 10µM) (n=3 per group). Data are shown as the mean ± SD. *P<0.05, **P<0.01, ***P<0.001, one-way ANOVA with Dunnett’s multiple comparisons test. (D) Representative images of myotube formation of C2C12 cells in myogenic differentiation media for 7 days in control conditions, or with BYL719 (2 or 10µM). Scale bar represents 200 μm.

## References

1. Agarwal S, Loder SJ, Breuler C, Li J, Cholok D, Brownley C, Peterson J, Hsieh HH, Drake J, Ranganathan K, Niknafs YS, Xiao W, Li S, Kumar R, Tompkins R, Longaker MT, Davis TA, Yu PB, Mishina Y, Levi B. 2017. Strategic Targeting of Multiple BMP Receptors Prevents Trauma-Induced Heterotopic Ossification. Molecular Therapy 25:1974–1987. doi:10.1016/j.ymthe.2017.01.008

2. Agnew C, Ayaz P, Kashima R, Loving HS, Ghatpande P, Kung JE, Underbakke ES, Shan Y, Shaw DE, Hata A, Jura N. 2021. Structural basis for ALK2/BMPR2 receptor complex signaling through kinase domain oligomerization. Nat Commun 12. doi:10.1038/s41467-021-25248-5

3. Alessi Wolken DM, Idone V, Hatsell SJ, Yu PB, Economides AN. 2017. The obligatory role of Activin A in the formation of heterotopic bone in Fibrodysplasia Ossificans Progressiva. Bone 109:210–217. doi:10.1016/j.bone.2017.06.011

4. Aykul S, Huang L, Wang L, Das NM, Reisman S, Ray Y, Zhang Q, Rothman N, Nannuru KC, Kamat V, Brydges S, Troncone L, Johnsen L, Yu PB, Fazio S, Lees-Shepard J, Schutz K, Murphy AJ, Economides AN, Idone V, Hatsell SJ. 2022. Anti-ACVR1 antibodies exacerbate heterotopic ossification in fibrodysplasia ossificans progressiva (FOP) by activating FOP-mutant ACVR1. Journal of Clinical Investigation 132. doi:10.1172/JCI153792

5. Barruet E, Morales BM, Cain CJ, Ton AN, Wentworth KL, Chan T V., Moody TA, Haks MC, Ottenhoff TH, Hellman J, Nakamura MC, Hsiao EC. 2018. NF-κB/MAPK activation underlies ACVR1-mediated inflammation in human heterotopic ossification. JCI Insight 3. doi:10.1172/jci.insight.122958

6. Bravenboer N, Micha D, Triffit JT, Bullock AN, Ravazollo R, Bocciardi R, Di Rocco M, Netelenbos JC, Ten Dijke P, Sánchez-Duffhues G, Kaplan FS, Shore EM, Pignolo RJ, Seemann P, Ventura F, Beaujat G, Eekhoff EMW, Pals G. 2015. Clinical Utility Gene Card for: Fibrodysplasia ossificans progressiva. European Journal of Human Genetics 23. doi:10.1038/ejhg.2014.274

7. Cardoso F, Paluch-Shimon S, Senkus E, Curigliano G, Aapro MS, André F, Barrios CH, Bergh J, Bhattacharyya GS, Biganzoli L, Boyle F, Cardoso M-J, Carey LA, Cortés J, el Saghir NS, Elzayat M, Eniu A, Fallowfield L, Francis PA, Gelmon K, Gligorov J, Haidinger R, Harbeck N, Hu X, Kaufman B, Kaur R, Kiely BE, Kim S-B, Lin NU, Mertz SA, Neciosup S, Offersen B v, Ohno S, Pagani O, Prat A, Penault-Llorca F, Rugo HS, Sledge GW, Thomssen C, Vorobiof DA, Wiseman T, Xu B, Norton L, Costa A, Winer EP. 2020. 5th ESO-ESMO international consensus guidelines for advanced breast cancer (ABC 5). Ann Oncol 31:1623–1649. doi:10.1016/j.annonc.2020.09.010

8. Chaikuad A, Alfano I, Kerr G, Sanvitale CE, Boergermann JH, Triffitt JT, Von Delft F, Knapp S, Knaus P, Bullock AN. 2012. Structure of the bone morphogenetic protein receptor ALK2 and implications for fibrodysplasia ossificans progressiva. Journal of Biological Chemistry 287:36990–36998. doi:10.1074/jbc.M112.365932

9. Chakkalakal SA, Uchibe K, Convente MR, Zhang D, Economides AN, Kaplan FS, Pacifici M, Iwamoto M, Shore EM. 2016. Palovarotene Inhibits Heterotopic Ossification and Maintains Limb Mobility and Growth in Mice With the Human ACVR1(R206H) Fibrodysplasia Ossificans Progressiva (FOP) Mutation. J Bone Miner Res 31:1666–75. doi:10.1002/jbmr.2820

10. Chakkalakal SA, Zhang D, Culbert AL, Convente MR, Caron RJ, Wright AC, Maidment A DA, Kaplan FS, Shore EM. 2012. An *Acvr1* R206H knock-in mouse has fibrodysplasia ossificans progressiva. Journal of Bone and Mineral Research. doi:10.1002/jbmr.1637

11. Convente MR, Chakkalakal SA, Yang E, Caron RJ, Zhang D, Kambayashi T, Kaplan FS, Shore EM. 2018. Depletion of Mast Cells and Macrophages Impairs Heterotopic Ossification in an Acvr1^R206H^Mouse Model of Fibrodysplasia Ossificans Progressiva. Journal of Bone and Mineral Research 33. doi:10.1002/jbmr.3304

12. Dey D, Bagarova J, Hatsell SJ, Armstrong KA, Huang L, Ermann J, Vonner AJ, Shen Y, Mohedas AH, Lee A, Eekhoff EMW, Van Schie A, Demay MB, Keller C, Wagers AJ, Economides AN, Yu PB. 2016. Two tissue-resident progenitor lineages drive distinct phenotypes of heterotopic ossification. Sci Transl Med 8. doi:10.1126/scitranslmed.aaf1090

13. Eisner C, Cummings M, Johnston G, Tung LW, Groppa E, Chang C, Rossi FMV. 2020. Murine Tissue-Resident PDGFRα+ Fibro-Adipogenic Progenitors Spontaneously Acquire Osteogenic Phenotype in an Altered Inflammatory Environment. Journal of Bone and Mineral Research 35:1525–1534. doi:10.1002/jbmr.4020

14. Ford-Hutchinson AF, Ali Z, Lines SE, Hallgrímsson B, Boyd SK, Jirik FR. 2007. Inactivation of Pten in osteo-chondroprogenitor cells leads to epiphyseal growth plate abnormalities and skeletal overgrowth. Journal of Bone and Mineral Research. doi:10.1359/jbmr.070420

15. Fujita T, Azuma Y, Fukuyama R, Hattori Y, Yoshida C, Koida M, Ogita K, Komori T. 2004. Runx2 induces osteoblast and chondrocyte differentiation and enhances their migration by coupling with PI3K-Akt signaling. Journal of Cell Biology. doi:10.1083/jcb.200401138

16. Fukuda T, Scott G, Komatsu Y, Araya R, Kawano M, Ray MK, Yamada M, Mishina Y. 2006. Generation of a mouse with conditionally activated signaling through the BMP receptor, ALK2. Genesis. doi:10.1002/dvg.20201

17. Furet P, Guagnano V, Fairhurst RA, Imbach-Weese P, Bruce I, Knapp M, Fritsch C, Blasco F, Blanz J, Aichholz R, Hamon J, Fabbro D, Caravatti G. 2013. Discovery of NVP-BYL719 a potent and selective phosphatidylinositol-3 kinase alpha inhibitor selected for clinical evaluation. Bioorg Med Chem Lett 23:3741–3748. doi:10.1016/j.bmcl.2013.05.007

18. Gámez Beatriz, Rodríguez□Carballo E, Graupera M, Rosa JL, Ventura F. 2016. Class I PI□3□Kinase Signaling Is Critical for Bone Formation Through Regulation of SMAD1 Activity in Osteoblasts. Journal of Bone and Mineral Research 31:1617–1630.

19. Gámez B., Rodríguez-Carballo E, Graupera M, Rosa JL, Ventura F. 2016. Class I PI-3-Kinase Signaling Is Critical for Bone Formation Through Regulation of SMAD1 Activity in Osteoblasts. Journal of Bone and Mineral Research 31. doi:10.1002/jbmr.2819

20. Groppe JC, Wu J, Shore EM, Kaplan FS. 2011. In vitro analyses of the dysregulated R206H ALK2 kinase-FKBP12 interaction associated with heterotopic ossification in FOPCells Tissues Organs. pp. 291–295. doi:10.1159/000324230

21. Hatsell SJ, Idone V, Wolken DMA, Huang L, Kim HJ, Wang L, Wen X, Nannuru KC, Jimenez J, Xie L, Das N, Makhoul G, Chernomorsky R, D’Ambrosio D, Corpina RA, Schoenherr CJ, Feeley K, Yu PB, Yancopoulos GD, Murphy AJ, Economides AN. 2015. *ACVR1 ^R206H^* receptor mutation causes fibrodysplasia ossificans progressiva by imparting responsiveness to activin A. Sci Transl Med 7:303ra137–303ra137. doi:10.1126/scitranslmed.aac4358

22. Hino K, Ikeya M, Horigome K, Matsumoto Y, Ebise H, Nishio M, Sekiguchi K, Shibata M, Nagata S, Matsuda S, Toguchida J. 2015. Neofunction of ACVR1 in fibrodysplasia ossificans progressiva. Proceedings of the National Academy of Sciences 112:15438–15443. doi:10.1073/pnas.1510540112

23. Hwang CD, Pagani CA, Nunez JH, Cherief M, Qin Q, Gomez-Salazar M, Kadaikal B, Kang H, Chowdary AR, Patel N, James AW, Levi B. 2022. Contemporary perspectives on heterotopic ossification. JCI Insight 7. doi:10.1172/jci.insight.158996

24. Hwang C, Pagani CA, Das N, Marini S, Huber AK, Xie LQ, Jimenez J, Brydges S, Lim WK, Nannuru KC, Murphy AJ, Economides AN, Hatsell SJ, Levi B. 2020. Activin A does not drive post-traumatic heterotopic ossification. Bone 138:115473. doi:10.1016/j.bone.2020.115473

25. Ikegami D, Akiyama H, Suzuki A, Nakamura T, Nakano T, Yoshikawa H, Tsumaki N. 2011. Sox9 sustains chondrocyte survival and hypertrophy in part through Pik3ca-Akt pathways. Development. doi:10.1242/dev.057802

26. Jamieson S, Flanagan JU, Kolekar S, Buchanan C, Kendall JD, Lee W-J, Rewcastle GW, Denny WA, Singh R, Dickson J, Baguley BC, Shepherd PR. 2011. A drug targeting only p110α can block phosphoinositide 3-kinase signalling and tumour growth in certain cell types. Biochemical Journal. doi:10.1042/BJ20110502

27. Kan L, Liu Y, McGuire TL, Berger DMP, Awatramani RB, Dymecki SM, Kessler JA. 2009. Dysregulation of Local Stem/Progenitor Cells as a Common Cellular Mechanism for Heterotopic Ossification. Stem Cells 27:150–156. doi:10.1634/stemcells.2008-0576

28. Kaplan FS, Zasloff MA, Kitterman JA, Shore EM, Hong CC, Rocke DM. 2010. Early mortality and cardiorespiratory failure in patients with fibrodysplasia ossificans progressiva. Journal of Bone and Joint Surgery - Series A 92. doi:10.2106/JBJS.I.00705

29. Kaplan FS, Zeitlin L, Dunn SP, Benor S, Hagin D, al Mukaddam M, Pignolo RJ. 2018. Acute and chronic rapamycin use in patients with Fibrodysplasia Ossificans Progressiva: A report of two cases. Bone 109:281–284. doi:10.1016/j.bone.2017.12.011

30. Lees-Shepard JB, Stoessel SJ, Chandler JT, Bouchard K, Bento P, Apuzzo LN, Devarakonda PM, Hunter JW, Goldhamer DJ. 2022. An anti-ACVR1 antibody exacerbates heterotopic ossification by fibro-adipogenic progenitors in fibrodysplasia ossificans progressiva mice. Journal of Clinical Investigation 132. doi:10.1172/JCI153795

31. Lees-Shepard JB, Yamamoto M, Biswas AA, Stoessel SJ, Nicholas SAE, Cogswell CA, Devarakonda PM, Schneider MJ, Cummins SM, Legendre NP, Yamamoto S, Kaartinen V, Hunter JW, Goldhamer DJ. 2018. Activin-dependent signaling in fibro/adipogenic progenitors causes fibrodysplasia ossificans progressiva. Nat Commun 9. doi:10.1038/s41467-018-02872-2

32. Madsen RR, Semple RK. 2022. PIK3CA-related overgrowth: silver bullets from the cancer arsenal? Trends Mol Med 28:255–257. doi:10.1016/j.molmed.2022.02.009

33. Matsuo K, Chavez RD, Barruet E, Hsiao EC. 2019. Inflammation in Fibrodysplasia Ossificans Progressiva and Other Forms of Heterotopic Ossification. Curr Osteoporos Rep 17:387–394. doi:10.1007/s11914-019-00541-x

34. Matsuo K, Lepinski A, Chavez RD, Barruet E, Pereira A, Moody TA, Ton AN, Sharma A, Hellman J, Tomoda K, Nakamura MC, Hsiao EC. 2021. ACVR1R206H extends inflammatory responses in human induced pluripotent stem cell-derived macrophages. Bone 153:116129. doi:10.1016/j.bone.2021.116129

35. Morin G, Degrugillier-Chopinet C, Vincent M, Fraissenon A, Aubert H, Chapelle C, Hoguin C, Dubos F, Catteau B, Petit F, Mezel A, Domanski O, Herbreteau G, Alesandrini M, Boddaert N, Boutry N, Broissand C, Han TK, Branle F, Sarnacki S, Blanc T, Guibaud L, Canaud G. 2022. Treatment of two infants with PIK3CA-related overgrowth spectrum by alpelisib. J Exp Med 219. doi:10.1084/jem.20212148

36. Nakashima K, Zhou X, Kunkel G, Zhang Z, Deng JM, Behringer RR, De Crombrugghe B. 2002. The novel zinc finger-containing transcription factor Osterix is required for osteoblast differentiation and bone formation. Cell. doi:10.1016/S0092-8674(01)00622-5

37. Patel NK, Huber AK, Levi B. 2022. Macrophage TGFβ signaling is critical for wound healing with heterotopic ossification after trauma.

38. Pignolo RJ, Bedford-Gay C, Liljesthröm M, Durbin-Johnson BP, Shore EM, Rocke DM, Kaplan FS. 2016. The Natural History of Flare-Ups in Fibrodysplasia Ossificans Progressiva (FOP): A Comprehensive Global Assessment. Journal of Bone and Mineral Research 31:650–656. doi:10.1002/jbmr.2728

39. Pignolo RJ, Shore EM, Kaplan FS. 2013. Fibrodysplasia ossificans progressiva: diagnosis, management, and therapeutic horizons. Pediatr Endocrinol Rev 10 **Suppl 2**:437–448.

40. Ramachandran A, Mehić M, Wasim L, Malinova D, Gori I, Blaszczyk BK, Carvalho DM, Shore EM, Jones C, Hyvönen M, Tolar P, Hill CS. 2021. Pathogenic ACVR1 R206H activation by Activin A□induced receptor clustering and autophosphorylation . EMBO J 40. doi:10.15252/embj.2020106317

41. Rosen CJ, Lees-Shepard JB, Nicholas S-AE, Stoessel SJ, Devarakonda PM, Schneider MJ, Yamamoto M, Goldhamer DJ. 2018. Palovarotene reduces heterotopic ossification in juvenile FOP mice but exhibits pronounced skeletal toxicity. doi:10.7554/eLife.40814.001

42. Sánchez-Duffhues G, Williams E, Benderitter P, Orlova V, van Wijhe M, Garcia de Vinuesa A, Kerr G, Caradec J, Lodder K, de Boer HC, Goumans MJ, Eekhoff EMW, Morales-Piga A, Bachiller-Corral J, Koolwijk P, Bullock AN, Hoflack J, ten Dijke P. 2019. Development of Macrocycle Kinase Inhibitors for ALK2 Using Fibrodysplasia Ossificans Progressiva-Derived Endothelial Cells. JBMR Plus 3. doi:10.1002/jbm4.10230

43. Shimono K, Tung W, Macolino C, Chi AH-T, Didizian JH, Mundy C, Chandraratna RA, Mishina Y, Enomoto-Iwamoto M, Pacifici M, Iwamoto M. 2011. Potent inhibition of heterotopic ossification by nuclear retinoic acid receptor-γ agonists. Nat Med. doi:10.1038/nm.2334

44. Shore EM, Xu M, Feldman GJ, Fenstermacher DA, Brown MA, Kaplan FS. 2006. A recurrent mutation in the BMP type I receptor ACVR1 causes inherited and sporadic fibrodysplasia ossificans progressiva. Nat Genet 38:525–527. doi:10.1038/ng1783

45. Sierra-Filardi E, Nieto C, Domínguez-Soto Á, Barroso R, Sánchez-Mateos P, Puig-Kroger A, López-Bravo M, Joven J, Ardavín C, Rodríguez-Fernández JL, Sánchez-Torres C, Mellado M, Corbí ÁL. 2014. CCL2 Shapes Macrophage Polarization by GM-CSF and M-CSF: Identification of CCL2/CCR2-Dependent Gene Expression Profile. The Journal of Immunology 192:3858–3867. doi:10.4049/jimmunol.1302821

46. Soleimani M, Nadri S. 2009. A protocol for isolation and culture of mesenchymal stem cells from mouse bone marrow. Nat Protoc 4:102–106. doi:10.1038/nprot.2008.221

47. Sorkin M, Huber AK, Hwang C, Iv WFC, Menon R, Li J, Vasquez K, Pagani C, Patel N, Li S, Visser ND, Niknafs Y, Loder S, Scola M, Nycz D, Gallagher K, Mccauley LK, Xu J, James AW, Agarwal S, Kunkel S, Mishina Y, Levi B. 2020. by monocytes in a mouse model of aberrant. Nat Commun. doi:10.1038/s41467-019-14172-4

49. Stanley A, Tichy ED, Kocan J, Roberts DW, Shore EM, Mourkioti F. 2022. Dynamics of skeletal muscle-resident stem cells during myogenesis in fibrodysplasia ossificans progressiva. NPJ Regen Med 7:1–15. doi:10.1038/s41536-021-00201-8

50. Torossian F, Guerton B, Anginot A, Alexander KA, Desterke C, Soave S, Tseng HW, Arouche N, Boutin L, Kulina I, Salga M, Jose B, Pettit AR, Clay D, Rochet N, Vlachos E, Genet G, Debaud C, Denormandie P, Genet F, Sims NA, Banzet S, Levesque JP, Lataillade JJ, Le Bousse-Kerdilès MC. 2017. Macrophage-derived oncostatin M contributes to human and mouse neurogenic heterotopic ossifications. JCI Insight 2. doi:10.1172/jci.insight.96034

51. Towler OW, Shore EM. 2022. BMP signaling and skeletal development in fibrodysplasia ossificans progressiva (FOP). Developmental Dynamics 251:164–177. doi:10.1002/dvdy.387

52. Tu B, Li J, Sun Z, Zhang T, Liu H, Yuan F, Fan C. 2022. Macrophage-Derived TGF-β and VEGF Promote the Progression of Trauma-Induced Heterotopic Ossification. Inflammation. doi:10.1007/s10753-022-01723-z

53. Upadhyay J, Xie LQ, Huang L, Das N, Stewart RC, Lyon MC, Palmer K, Rajamani S, Graul C, Lobo M, Wellman TJ, Soares EJ, Silva MD, Hesterman J, Wang L, Wen X, Qian X, Nannuru K, Idone V, Murphy AJ, Economides AN, Hatsell SJ. 2017. The Expansion of Heterotopic Bone in Fibrodysplasia Ossificans Progressiva Is Activin A-Dependent. Journal of Bone and Mineral Research 32:2489–2499. doi:10.1002/jbmr.3235

54. Valer JA, Sánchez□de□Diego C, Gámez B, Mishina Y, Rosa JL, Ventura F. 2019a. Inhibition of phosphatidylinositol 3□kinase α (PI 3Kα) prevents heterotopic ossification . EMBO Mol Med. doi:10.15252/emmm.201910567

55. Valer JA, Sánchez□de□Diego C, Gámez B, Mishina Y, Rosa JL, Ventura F. 2019b. Inhibition of phosphatidylinositol 3□kinase α (PI3Kα) prevents heterotopic ossification. EMBO Mol Med 11. doi:10.15252/emmm.201910567

56. Van Dinther M, Visser N, De Gorter DJJ, Doorn J, Goumans MJ, De Boer J, Ten Dijke P. 2010. ALK2 R206H mutation linked to fibrodysplasia ossificans progressiva confers constitutive activity to the BMP type I receptor and sensitizes mesenchymal cells to BMP-induced osteoblast differentiation and bone formation. Journal of Bone and Mineral Research 25:1208–1215. doi:10.1002/jbmr.091110

57. Wang JH, Liu YZ, Yin LJ, Chen L, Huang J, Liu Y, Zhang RX, Zhou LY, Yang QJ, Luo JY, Zuo G wei, Deng ZL, He BC. 2013. BMP9 and COX-2 form an important regulatory loop in BMP9-induced osteogenic differentiation of mesenchymal stem cells. Bone 57:311–321. doi:10.1016/j.bone.2013.08.015

58. Wang X, Li F, Xie L, Crane J, Zhen G, Mishina Y, Deng R, Gao B, Chen H, Liu S, Yang P, Gao M, Tu M, Wang Y, Wan M, Fan C, Cao X. 2018. Inhibition of overactive TGF-β attenuates progression of heterotopic ossification in mice. Nat Commun 9. doi:10.1038/s41467-018-02988-5

59. Wang X, Li F, Xie L, Crane J, Zhen G, Mishina Y, Deng R, Gao B, Chen H, Liu S, Yang P, Gao M, Tu M, Wang Y, Wan M, Fan C, Cao X. n.d. progression of heterotopic ossi fi cation in mice. Nat Commun. doi:10.1038/s41467-018-02988-5

60. Yu PB, Deng DY, Lai CS, Hong CC, Cuny GD, Bouxsein ML, Hong DW, McManus PM, Katagiri T, Sachidanandan C, Kamiya N, Fukuda T, Mishina Y, Peterson RT, Bloch KD. 2008. BMP type I receptor inhibition reduces heterotopic ossification. Nat Med. doi:10.1038/nm.1888

