## Supplementary figures and images for "PI3Kα inhibition blocks osteochondroprogenitor specification and the hyper-inflammatory response to prevent heterotopic ossification"

### Figure8-Figure Supplement 1

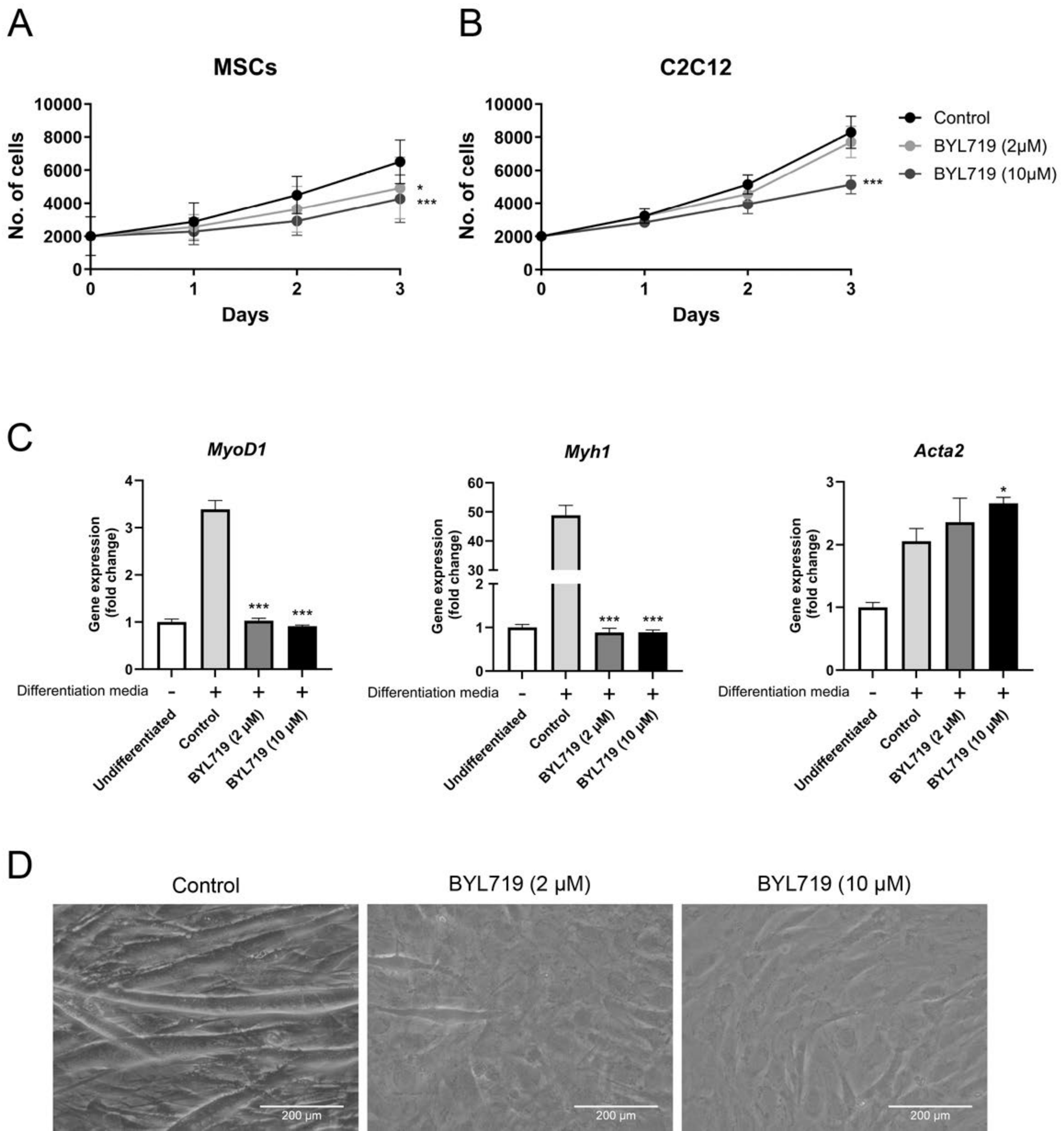

### Figure 1- Figure supplement 1

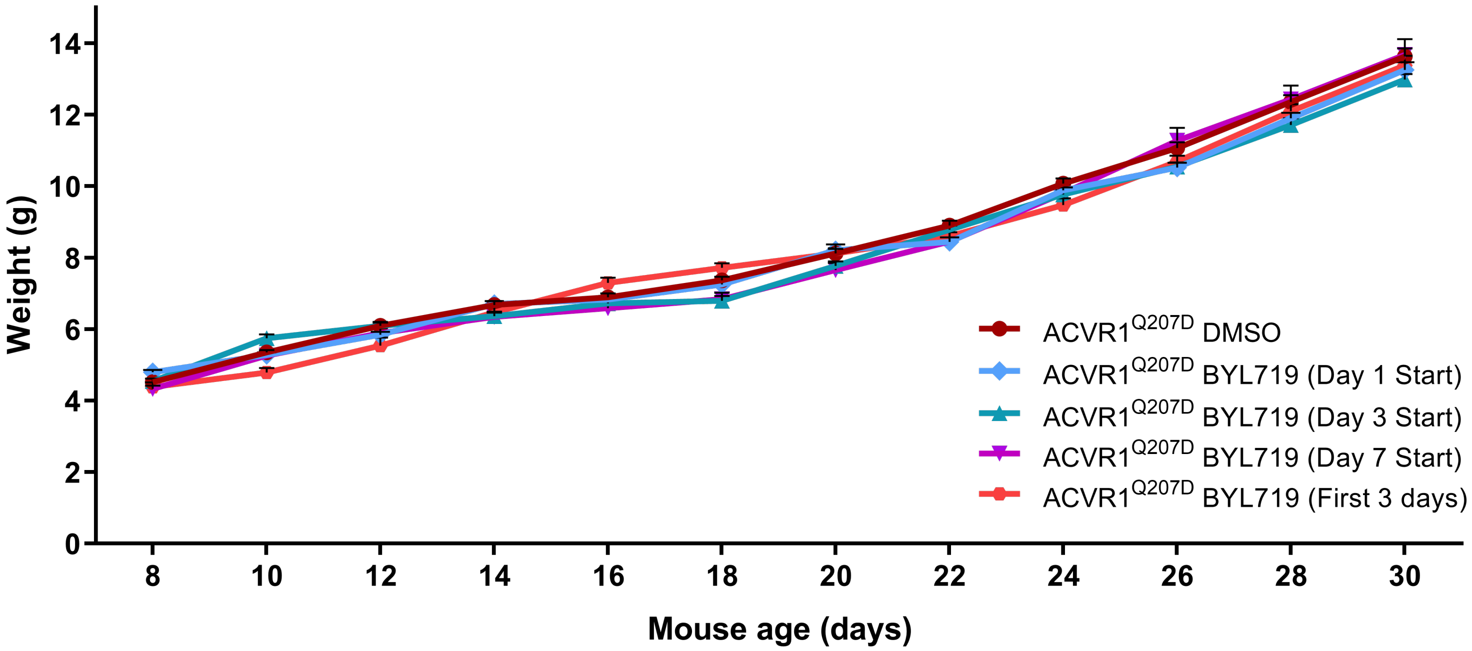

### Figure 2- Figure supplement 1

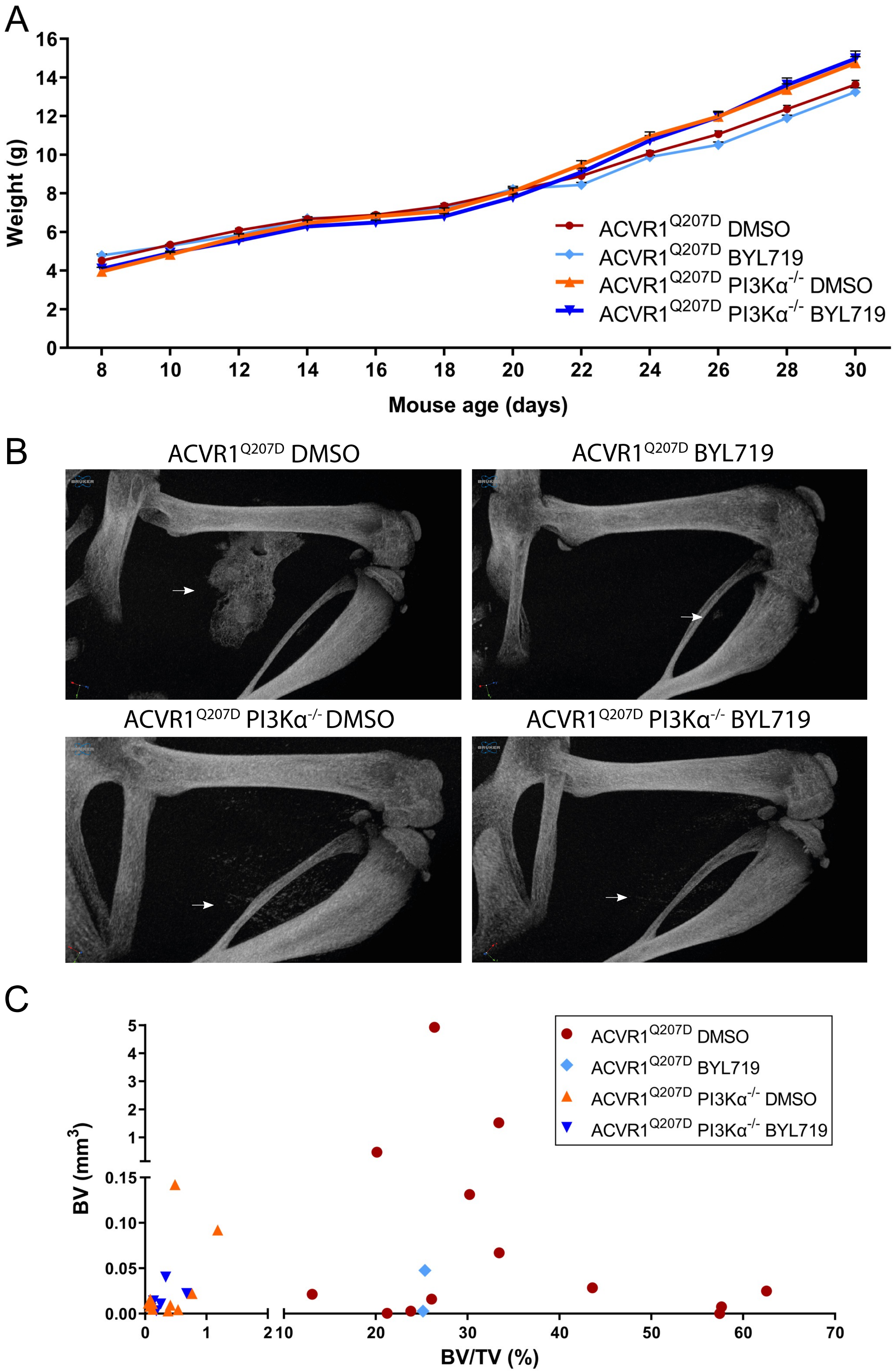

### Figure 2- Figure supplement 2

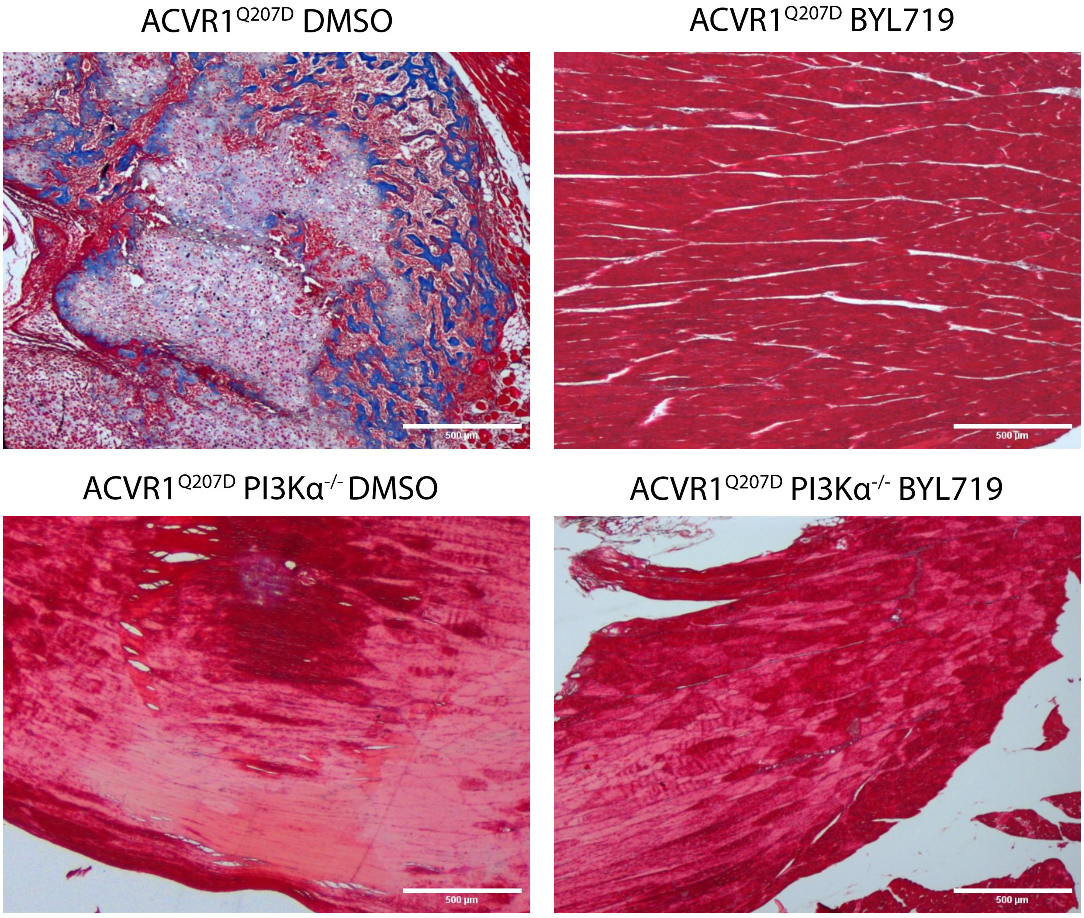

### Figure 3- Figure supplement 1

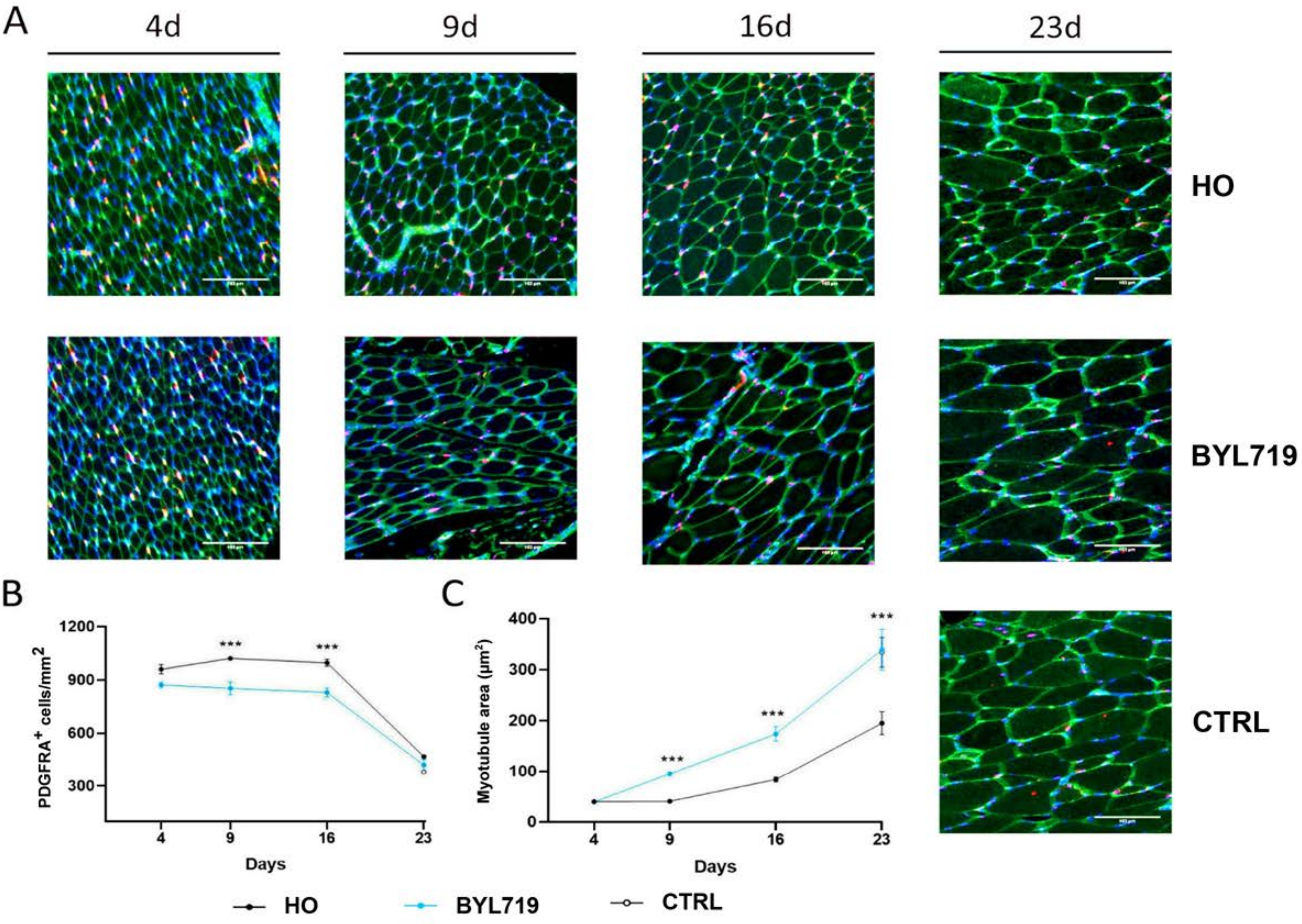

### Figure 4- Figure supplement 1

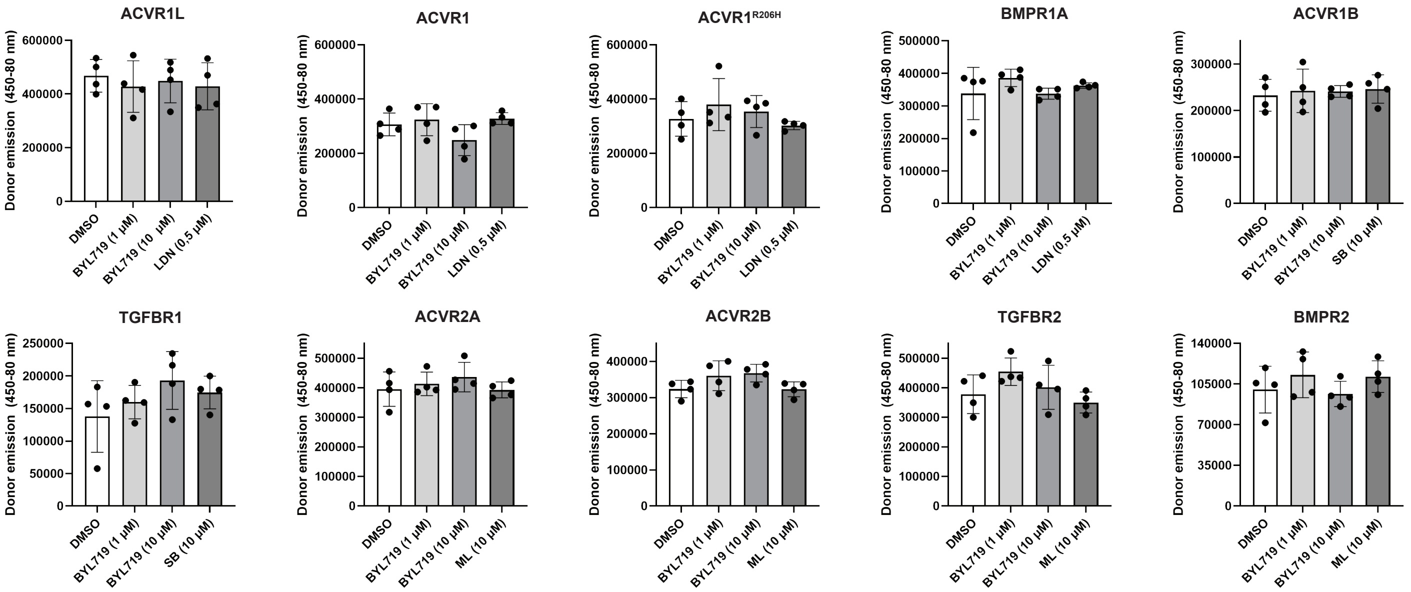

### Figure 5- Figure supplement 1

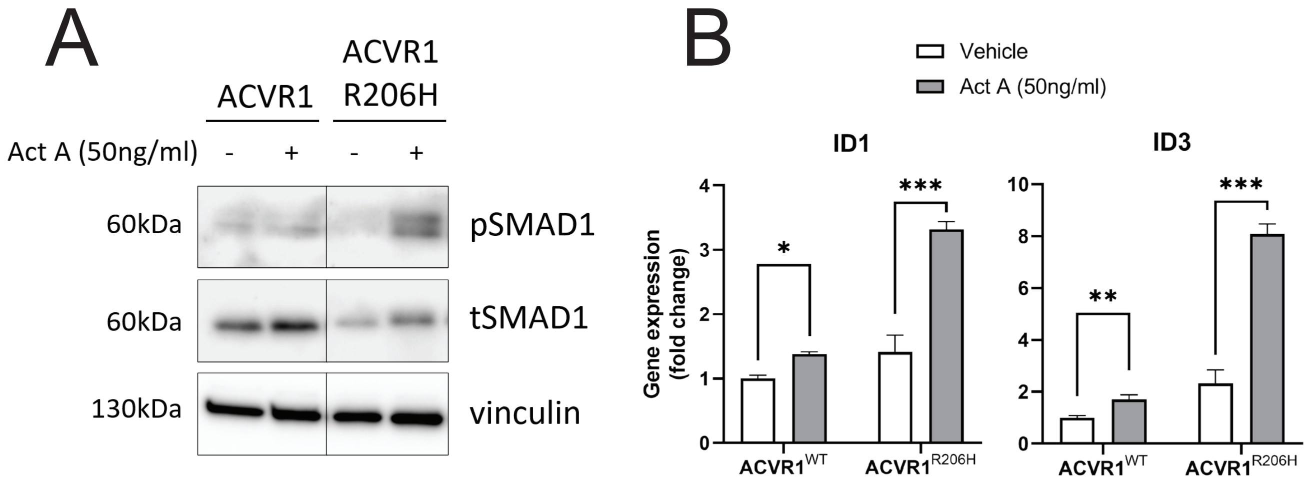

### Figure 6- Figure supplement 1

A

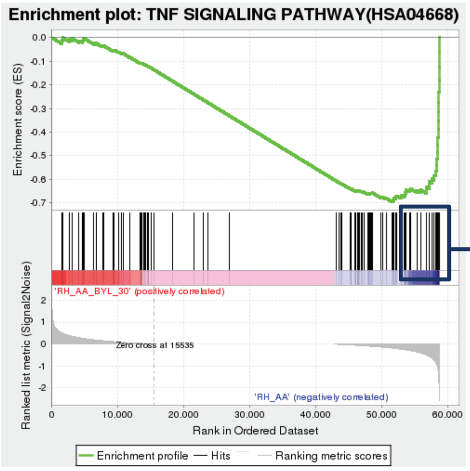


ES= -0.70, NES= -1.51, FDR= 0.13

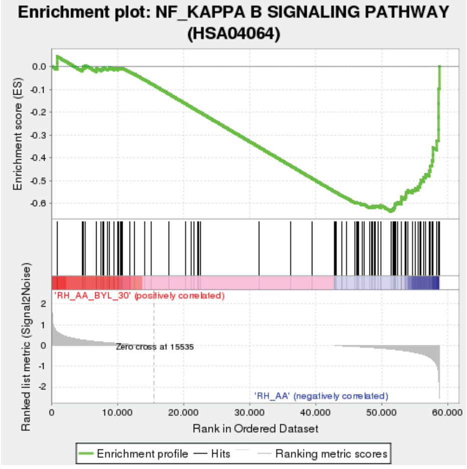


ES= -0.63, NES= -1.54, FDR= 0.18

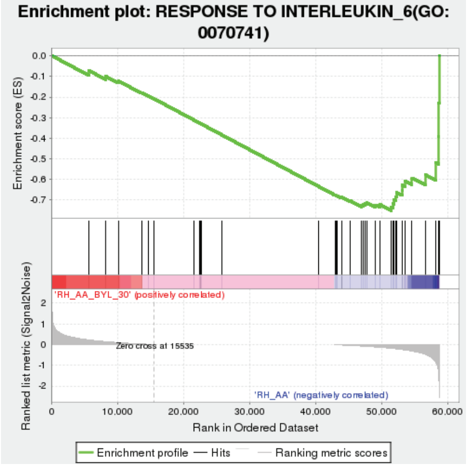


ES= -0.75, NES= -1.55, FDR= 0.24

B

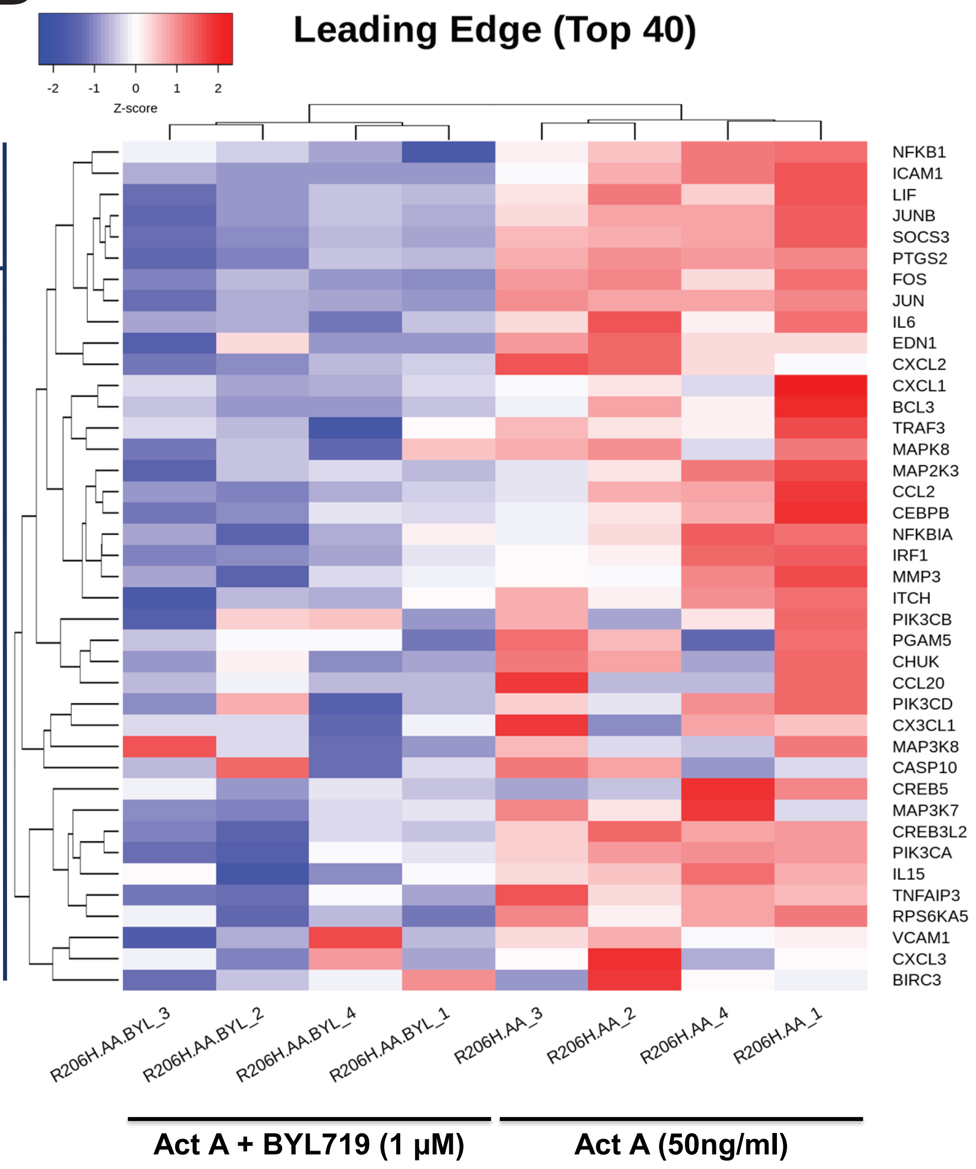

### Figure 7- Figure supplement 1

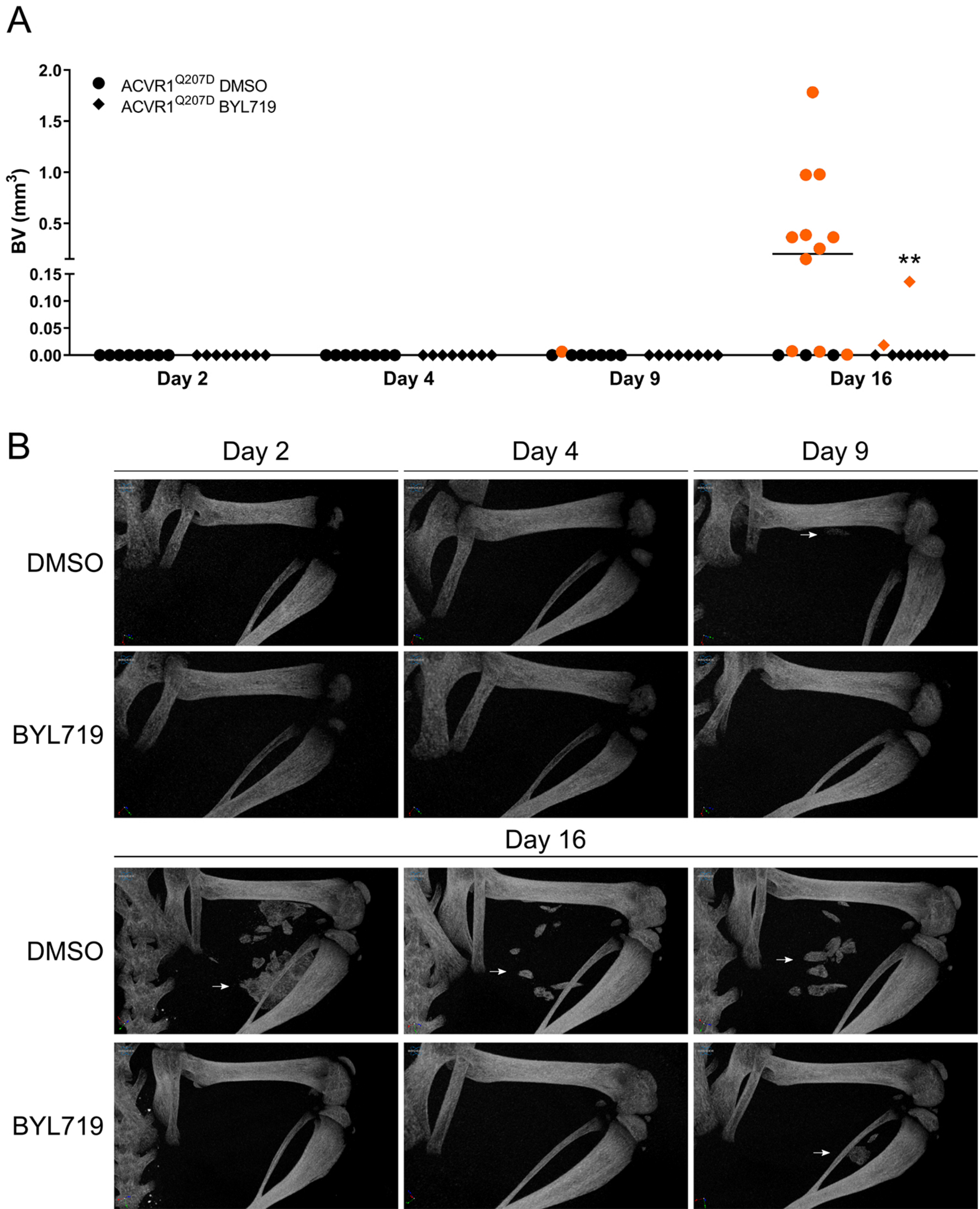

### Figure 7- Figure supplement 2

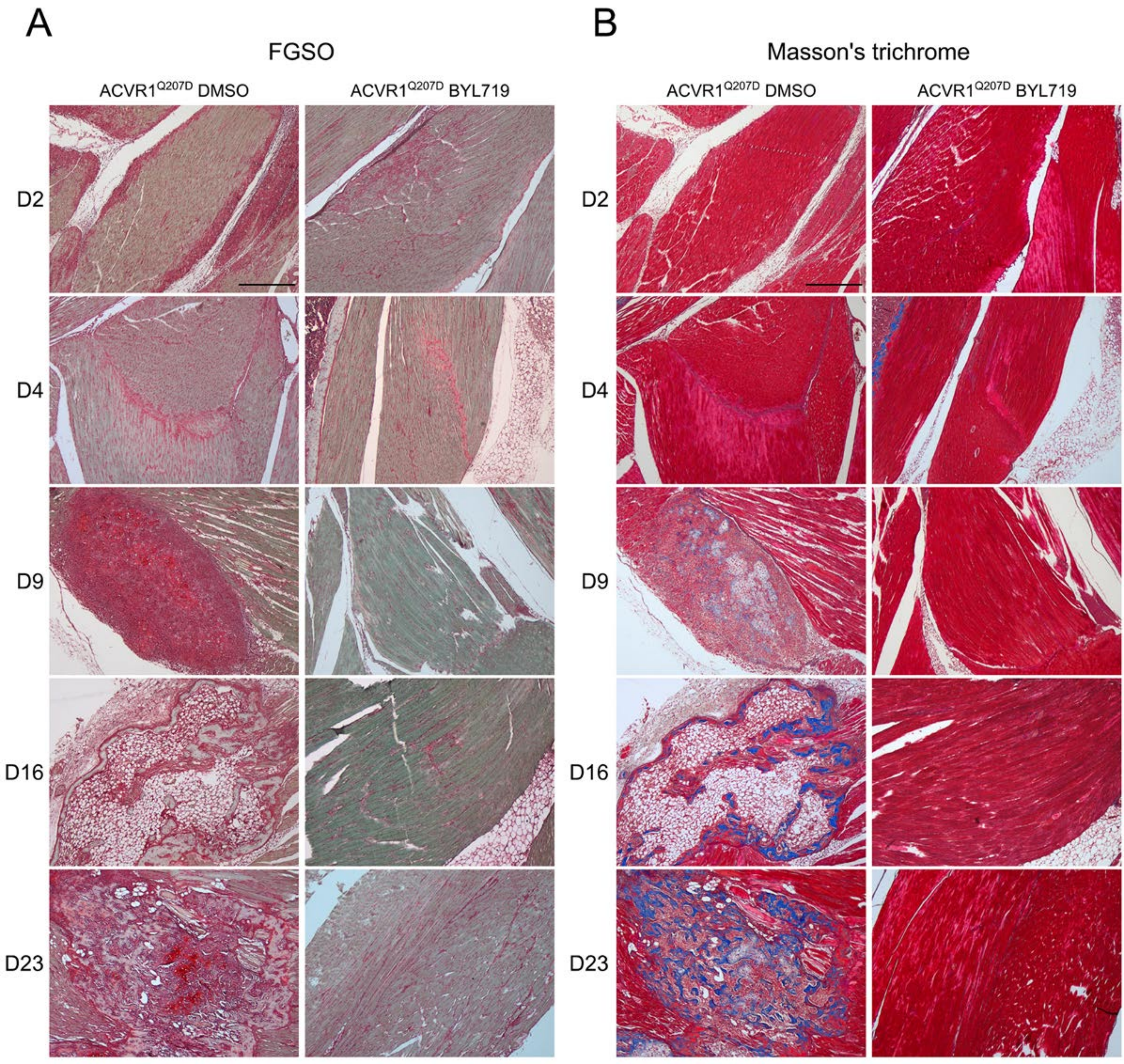
